# Lung, spleen and oesophagus tissue remains stable for scRNAseq in cold preservation

**DOI:** 10.1101/741405

**Authors:** E. Madissoon, A. Wilbrey-Clark, R.J. Miragaia, K. Saeb-Parsy, K. Mahbubani, N. Georgakopoulos, P. Harding, K. Polanski, K. Nowicki-Osuch, R.C. Fitzgerald, K.W. Loudon, J.R. Ferdinand, M.R Clatworthy, A. Tsingene, S. Van Dongen, M. Dabrowska, M. Patel, M.J.T. Stubbington, S. Teichmann, O. Stegle, K.B. Meyer

**Affiliations:** Wellcome Sanger Institute, Cambridge, UK; The European Bioinformatics Institute (EMBL-EBI), Wellcome Genome Campus, Hinxton, Cambridgeshire, CB10 1SD; Department of Surgery, University of Cambridge and NIHR Cambridge Biomedical Research Centre, Cambridge, CB2 0QQ; MRC Cancer Unit, Hutchinson-MRC Research Centre, University of Cambridge, Cambridge, CB2 0XZ; Molecular Immunology Unit, Department of Medicine, Cambridge, CB2 0QQ; 10x Genomics Inc., 6230 Stoneridge Mall Road, Pleasanton, CA 94588, USA

**Keywords:** Single cell RNA sequencing, human, spleen, oesophagus, lung, ischaemic time

## Abstract

**Background:** The Human Cell Atlas is a large international collaborative effort to map all cell types of the human body. Single cell RNA sequencing can generate high quality data for the delivery of such an atlas. However, delays between fresh sample collection and processing may lead to poor data and difficulties in experimental design. Despite this, there has not yet been a systematic assessment of the effect of cold storage time on the quality of scRNAseq

**Results:** This study assessed the effect of cold storage on fresh healthy spleen, oesophagus and lung from ≥5 donors over 72 hours. We collected 240,000 high quality single cell transcriptomes with detailed cell type annotations and whole genome sequences of donors, enabling future eQTL studies. Our data provide a valuable resource for the study of these three organs and will allow cross-organ comparison of cell types.

We see little effect of cold ischaemic time on cell viability, yield, total number of reads per cell and other quality control metrics in any of the tissues within the first 24 hours. However, we observed higher percentage of mitochondrial reads, indicative of cellular stress, and increased contamination by background “ambient RNA” reads in the 72h samples in spleen, which is cell type specific.

**Conclusions:** In conclusion, we present robust protocols for tissue preservation for up to 24 hours prior to scRNAseq analysis. This greatly facilitates the logistics of sample collection for Human Cell Atlas or clinical studies since it increases the time frames for sample processing.

## Background

High-throughput single cell RNA sequencing (scRNAseq) techniques have developed rapidly in recent years, making it feasible to generate transcriptional profiles from thousands of cells in parallel (1,2,3,4,5,6,7). This technology has deepened our understanding of the cell types within tissues, their interactions and cellular states (1,8,9,10,4,10,11,12,13,14,15,16). It is also a cornerstone of the Human Cell Atlas Project (HCA; 17,18,19), a large collaborative initiative which aims to identify every cell type in the human body. Human samples present particular logistical challenges: the clinic may be distant from the processing lab and tissue can become available at short-notice and / or at inconvenient times. These scenarios necessitate a fast, simple method of preserving samples that requires minimal processing at the clinic.

To address the logistical difficulties and rapid transcriptional changes / stress responses observed upon tissue dissociation (20) or storage (21), a range of cell freezing or fixation methods have been developed. Guillaumet-Adkins et al (2017, 22) demonstrate that although viability is reduced, the transcriptional profiles from cultured cells or minced mouse tissue biopsies cryopreserved with DMSO are not significantly altered. However, some cell types are more vulnerable to freezing than others, for example in human endometrium biopsies stromal cells survive freezing better than epithelial cells (23). Fixation of cells with traditional cross-linking fixatives (24), reversible-crosslinkers (25), non-crosslinking alternatives such as methanol (26) and other novel stabilization reagents (27) has also been tested. Fixation halts transcriptional change and stabilizes cell types, although it usually creates 3’ bias. Thus far these agents have been tested on dissociated cells, or at best minced tissues, rather than intact tissue pieces. Unfortunately dissociation prior to transportation is often not practical with human clinical samples and dissociating preserved / fixed tissue pieces using traditional mechanical or enzymatic dissociation methods is often challenging.

Hypothermic preservation of intact tissues, as used during organ transplant procedures, has been optimized to reduce the effects of ischaemia (lack of blood supply) and hypoxia (oxygen deficiency) during storage at 4°C (28). Clinically, kidneys are transplanted with a median cold ischaemic time of 13 hours and maximum around 35h; lungs with median 6.4h and maximum 14h. However, human kidney and pancreas maintain their function even after 72h storage in University of Wisconsin solution, and liver for up to 30h (29). Wang et al (30) demonstrated that intact mouse kidneys could be stored in HypoThermasol FRS media for up to 72h before dissociation and scRNAseq without altering the transcriptomic profile or cellular heterogeneity of kidney-resident immune cells. Considering human tissue research, this method has major advantages. Firstly, it requires no processing of the sample at the collection site; the clinician can immerse an intact piece of tissue in cold Hypothermasol FRS solution and store or ship this on ice to the receiving laboratory, where all other tissue processing can take place. This can be done in a standardized and reproducible way. Secondly, it utilizes a commercially-available, chemically-defined, non-toxic and ready-to-use hypothermic preservation solution, designed to mimic clinical organ preservation.

One limitation of the Wang et al study, however, was that it only studied murine kidney. To provide maximum utility for human research, scRNAseq from multiple human organs with different ischaemic sensitivities is required. Ferreira et al (21) saw organ-related variation in the number of genes that changed expression with post-mortem interval (warm ischaemia) in GTEx bulk RNA seq (31). For example, spleen showed relatively little change, whereas oesophagus mucosa and lung altered their transcriptional profiles more significantly; oesophagus showing a response that peaked and declined, whereas lung had a more sustained gene expression change. GTEx data (31) also demonstrates non-random, transcript-dependent changes in post-mortem RNA degradation and apparent gene expression (32,33).

In this study, we aimed to identify a method of tissue preservation that would stabilize intact human tissue samples for scRNAseq but that requires minimal processing at the clinic and allows sample transportation time. In order to contribute to the Human Cell Atlas, we tested the method on three human primary tissues expected to have different sensitivities to ischaemia (21): spleen (most stable), oesophagus mucosa, and lung (least stable) (21). These tissues contain cell types ranging from immune cells to keratinocytes. Samples were obtained from deceased organ donors, rapidly perfused with cold organ preservation solution following death. Our dataset of 240,000 single cells includes the largest published datasets on human oesophagus and spleen to date, which we provide in an easy to browse data portal: www.tissuestabilitycellatlas.org. We show that storing intact tissue pieces from these 3 organs at 4°C in HypoThermosol FRS for 24h, or in most cases 72h, had little effect on the transcriptomic profile as determined by bulk and 10x Genomics 3’ single cell RNA sequencing. The diversity of populations observed in scRNAseq data was maintained over time. This protocol should be easily adopted by many clinical sites and permits at least a 24h time window for shipping of samples to collaborators, therefore increasing accessibility to fresh human tissue for research.

## Results and discussion

### Good scRNAseq data quality after cold storage

We obtained lung, oesophagus and spleen samples from 12 organ donors (Supplementary Table 1). The transplant surgeon assessed each organ as overall healthy in appearance. Whole genome sequencing (WGS) was carried out for each individual, confirming that none of the study participants displayed gross genomic abnormalities. Furthermore, for each donor, histology sections were produced from the 12 or 24h time points of each tissue, stained with hematoxylin and eosin and assessed by a pathologist (Supplementary Figure 1). This confirmed all tissue sections as healthy, except one donor with possible lung hypertension. Heterogeneity between tissue sections, for example the presence of glands, and amount of inflammation in some sections (Supplementary Figure 1), is likely to impact profiling by scRNAseq.

Samples of lung parenchyma, oesophagus mid-region and spleen (n≥5; experimental design, Figure 1a) were placed into 4°C HypoThermasol FRS solution immediately after collection (within 2 hours of cessation of circulation with cold perfusion) and were kept at 4°C until used for scRNAseq. For the majority of lung donors (n=5) tissue pieces were also flash frozen at the clinic, before transport to the processing site for bulk RNA sequencing.

**Figure 1:**
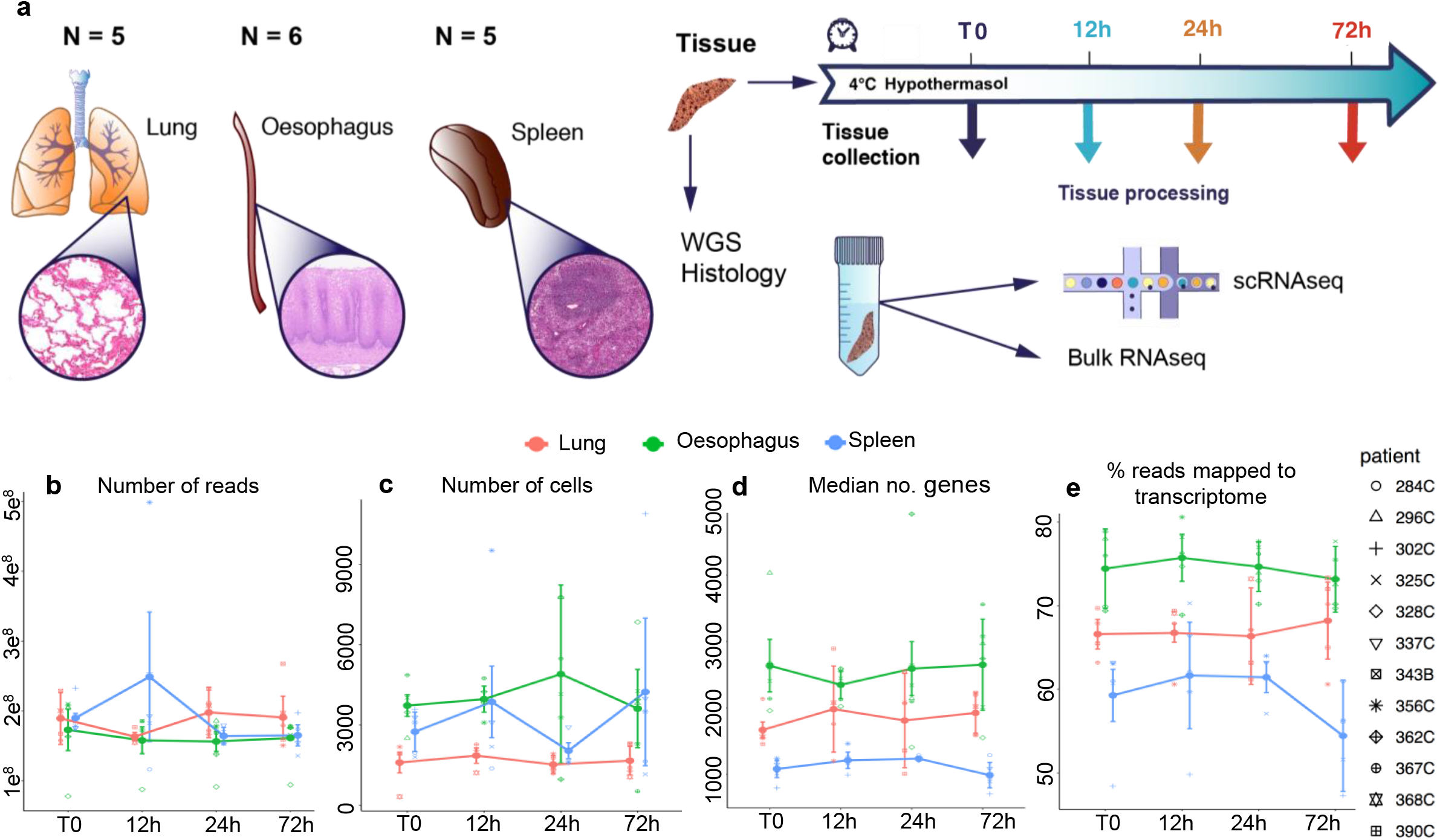
scRNA-seq quality metrics remain stable for at least 24 hours of cold storage. (a) Experimental design: Samples from lung, oesophagus and spleen were collected from 5 or 6 donors each and stored as whole organ pieces at 4°C for different time points prior to tissue processing for scRNAseq and bulk RNAseq. (b-e) Change of quality metrics of scRNA-seq data obtained with time, showing (b) number of reads per sample, (c) number of cells per sample, (d) median number of genes detected per cell and (e) the number of genes confidently mapped to the transcriptome.

Following transport, fresh samples were either dissociated immediately (T0) or stored at 4°C for 12, 24 or 72 hours cold ischaemic time prior to processing to single cell suspension (Figure 1b). While the T0 time point varied depending on the length of the organ transplant procedure, time required to collect samples in the clinic and speed of courier delivery (on average 4 hours of cold ischaemia from cessation of circulation to receipt of tissue at the processing laboratory), other time points were processed at 12h, 24h and 72h of ischaemic time. Cells were analysed by 10X 3’ v2 scRNA-seq (Figure 1c) and the number of cells obtained for each sample is given in Supplementary Table 2. At each time point, tissue pieces were also flash frozen for bulk RNA-seq analysis.

After alignment and normalisation of scRNA-seq data, quality control metrics were assessed for all samples (Figure 1, Supplementary Figure 2). The number of reads per sample, number of cells per sample, median number of genes per cell and other quality metrics did not change significantly over time for lung and oesophagus, but we did observe changes in spleen after 24h (Figure 1b-d, Supplementary Figure 2). The percentage of reads confidently (QC=255) mapped to the transcriptome was stable for all samples except for spleen at 72 hours (Figure 1.e). RNA quality was good and did not change with time in any of the tissues (Supplementary Table 3; RIN > 7 for the majority of bulk RNA seq samples, with the exception of four lower quality outliers in spleen mainly from a single donor). We conclude that in terms of quality metrics, we do not detect changes that are associated with the length of cold storage.

### Reduced scRNA-seq data quality by 72h in spleen

While the majority of quality values did not change with time, we did observe a decline in reads confidently mapped to the transcriptome for spleen. Further investigation showed a significantly decreased percentage of reads in exons (Figure 2a,b), and increased percentage of reads in introns (Figure 2c,d) in samples from the 72h time point in spleen (p-value <0.05) but not in other tissues or time points (Figure 2.b,d). This skewing between intronic and exonic reads becomes even more apparent when only the top and bottom quartile of cells (with respect to intronic and exonic alignment) are examined over time (Supplementary Figure 3). This result implies that non-spliced reads are more stable to degradation; presumably because nuclear mRNA is less susceptible to enzymatic degradation.

**Figure 2:**
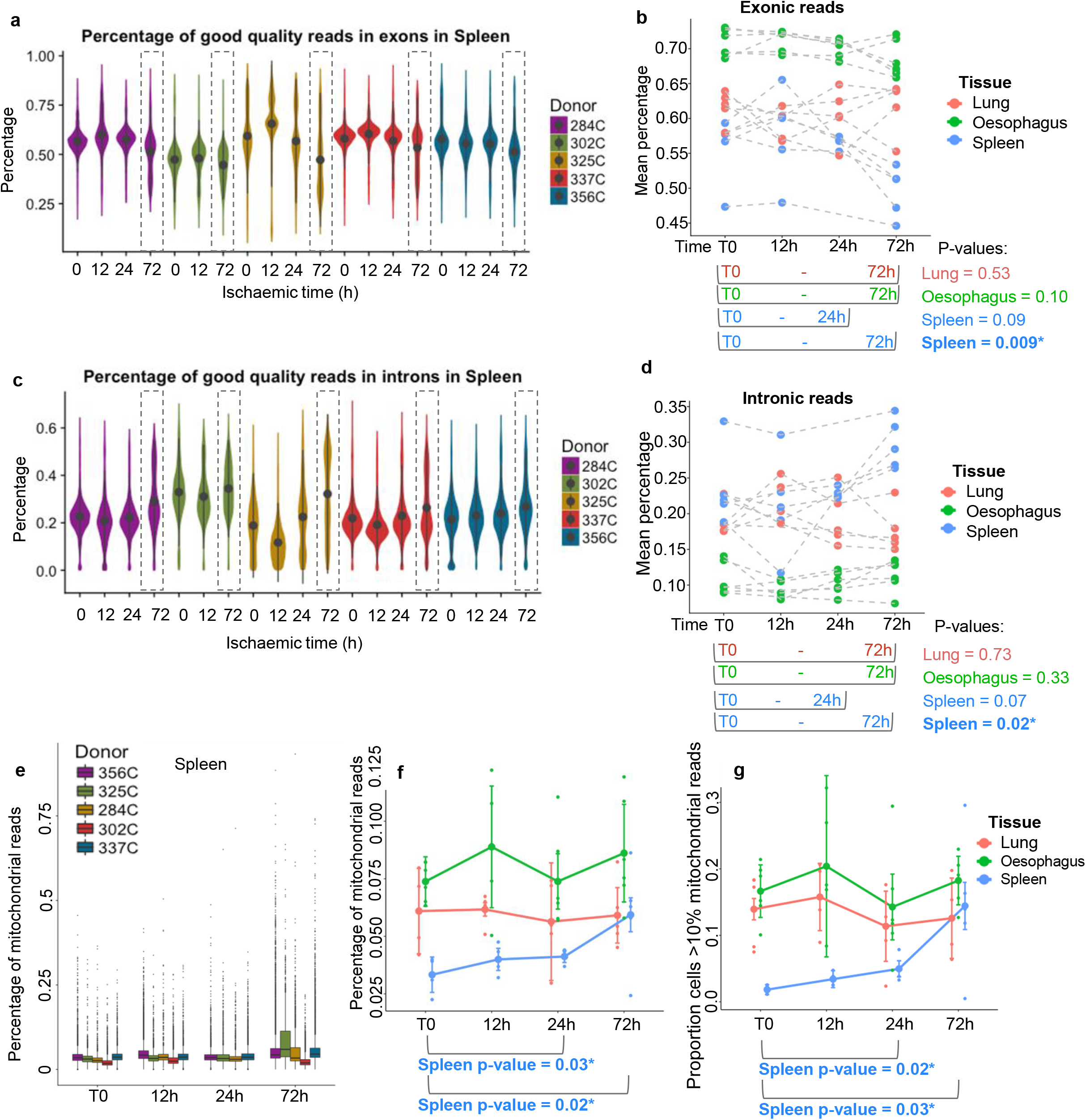
Exploration of loss of data quality with time in spleen compared to other organs. (a) Violin plot of good quality reads mapped to exons in spleen, (b) mean percentage of good quality exonic reads in all organs, (c) violin plot of good quality reads per exon in spleen, (d) mean percentage of intronic reads across all organs, (e) box plot of percentage of mitochondrial reads in spleen, (f) mean percentage of mitochondrial reads across all organs, and (e) percentage of cells with greater than 10% mitochondrial reads. The tissue of origin is indicated by colour.

We next assessed the proportion of mitochondrial reads. This is a commonly used quality metric indicative of cellular stress (34), for example induced by tissue dissociation or storage. Cells with high percentages of mitochondrial reads are generally excluded from analysis (35). In our data the fraction of mitochondrial reads was low, with no significant change in proportion, except in spleen where mitochondrial reads increase by 72h in 4 out of 5 donors (Figure 2e,f). This is also apparent when examining the number of cells with mitochondrial percentage higher than 10%, which significantly increases with time in spleen only (Figure 2g).

#### Effect of time on doublet rates and empty droplets

‘Scrublet’ (36) was used to calculate the doublet scores for each cell in every sample and these did not change with time for any of the three tissues (Supplementary Figure 4).

We next evaluated changes in non-cellular droplets (containing less than 100 reads). All droplet sequencing reactions generate many droplets that do not contain cells but capture acellular mRNA, often referred to as “ambient RNA” or “soup“(37). We define “ambient RNA” as 1-2 UMIs / droplet, “debris” as 3-100 UMIs / droplet and “cellular material” as >100 UMIs / droplet (Figure 3.a) to reflect the distribution of reads. The proportions of droplets containing UMIs in any of these intervals is not affected by time in spleen (Figure 3b), lung or oesophagus (Supplementary Figure 5a,b). However, the mean number of UMI increases in debris and decreases in cellular droplets by 72h (but not 24h) in spleen (Figure 3.c, d). This is not observed in lung or oesophagus, although mean debris UMIs are very variable in all three tissues.

**Figure 3:**
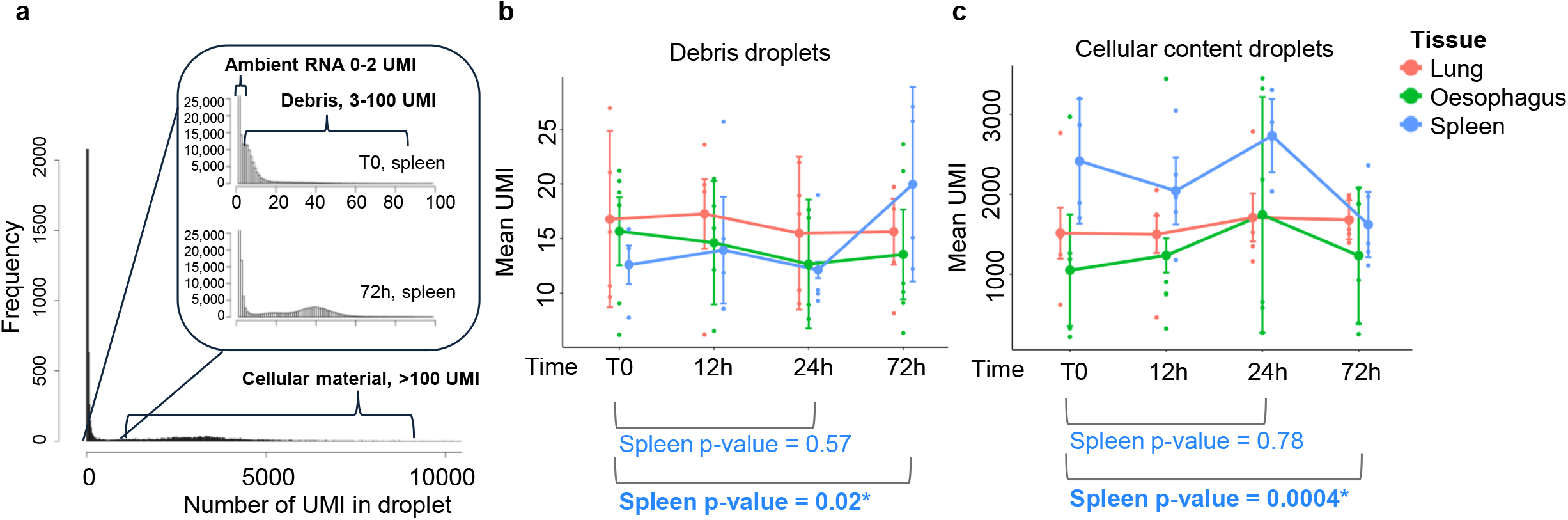
Loss of data quality is associated with increased “ambient RNA” and “debris” reads in the data. (a) Average spread of UMI counts per droplet in spleen, which were classified into ambient RNA, debris and cellular reads. (b) Proportions of droplets corresponding to each of these three categories. (c) Debris and cellular droplets with time across all tissues.

The increasing debris in spleen could indicate increased cellular death by 72h. Cell viability in spleen decreased between T0 and 72h in three of four recorded donors and lung viability decreased in all three of the recorded donors (Supplementary Figure 6). However, perhaps due to the small number of donors, this was not statistically significant. In addition, we observed significant variation in viability between samples that may be biological (donor variation) or technical (possibly due to samples being manually counted by multiple operators throughout the study).

#### Annotation of cell types

The gene expression count matrices from Cellranger output were used to perform sequential clustering of cells from either whole tissues or particular subclusters. The cell type identities of the clusters were determined and annotated by observation of expression of known cell type markers (Figure 4a-c, Supplementary Figure 7 a-c and Supplementary Table 2.)

**Figure 4:**
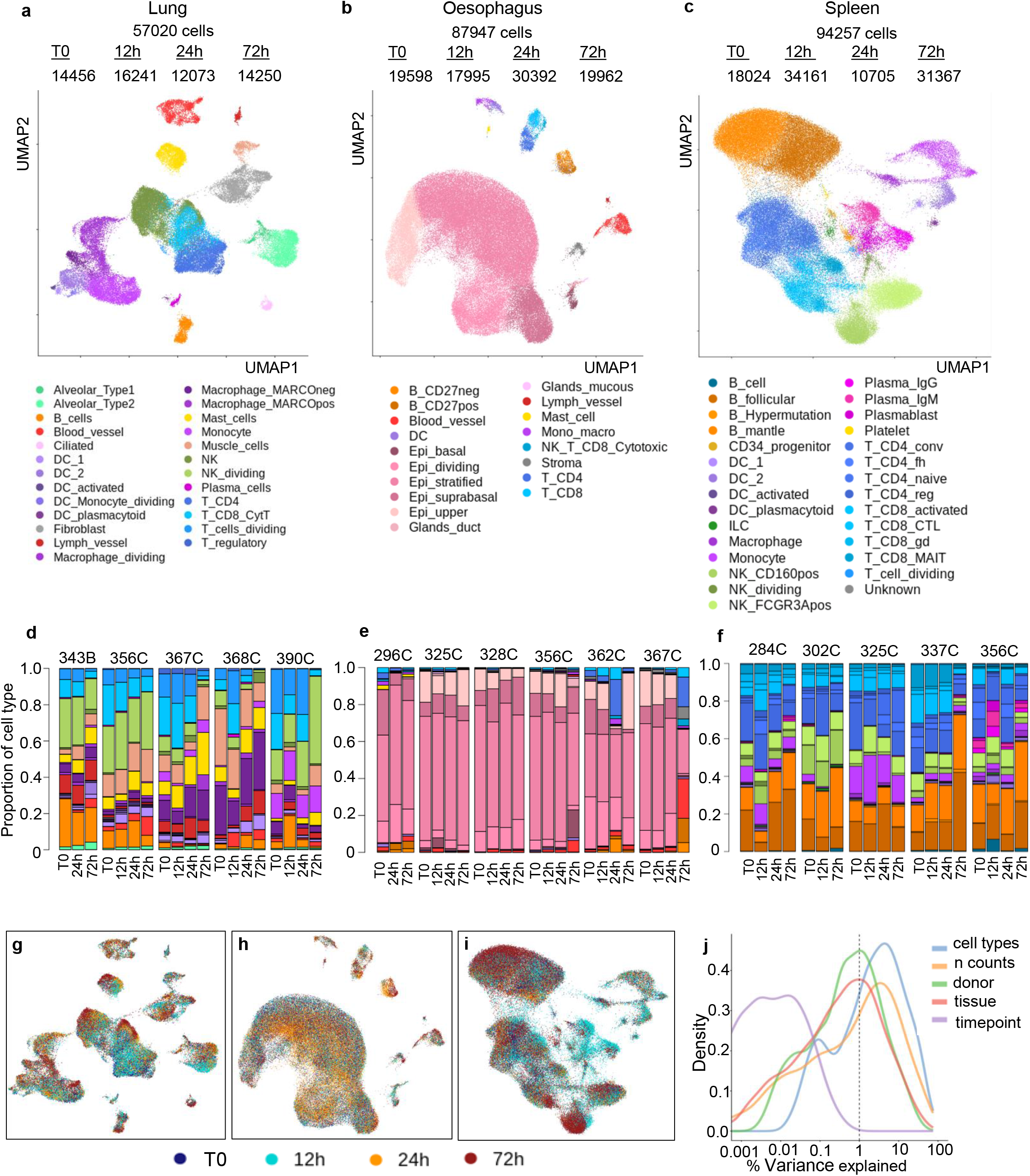
Cell types identified in different organs with time. (a) UMAP projections of scRNAseq data for lung (n= 57,020), (b) oesophagus (n= 87947) and (c) spleen (n= 94,257). (d-f) Proportions of cells identified per donor and per time point for (d) lung, (e) oesopaghus and (f) spleen. (g-h) Panels show the single cell UMAP plots for each organ with length of storage time highlighted. (j) Percent variance explained in the combined data set by cell types, n counts, donor, tissue and time points.

In lung, 57,020 cells passed quality control and represented 25 cell types. We detected ciliated, alveolar types 1 and 2 cells, as well as fibroblasts, muscle and endothelial cells both from blood and lymph vessels. The cell types identified from the immune compartment included NK, T and B-cells, as well as two types of macrophages, monocytes and dendritic cells (DC). Multiple DC populations such as conventional DC1, plasmacytoid DC (pcDC) and activated DC were detected and constituted 0.3% (163 cells), 0.08% (46 cells) and 0.2% (122 cells) of all cells, correspondingly. All donors contributed to every cluster. Dividing cells formed separate clusters for T-cells, DC, monocytes, NK and macrophages.

Oesophagus yielded 87,947 cells with over 90% belonging to the four major epithelial cell types: upper, stratified, suprabasal and dividing cells of the suprabasal layer. The additional cells from the basal layer of the epithelia clustered more closely to the gland duct and mucous secreting cells. While all donors contributed to the basal layer, only two samples from a total of 23 oesophagus samples provided the majority of the mucous secreting cells (0.06% from total; 55 cells; samples 325C, 12h and 356C, 72h). Nevertheless, each donor contributed to every cell type in all three tissues (Figure 4, Supplementary Figure 8 and Supplementary Table 2). Immune cells in the oesophagus include T-cells, B-cells, monocytes, macrophages, DCs and mast cells. Interestingly, almost 80% of the mast cells (87 cells) originated from a single donor (296C). Increased proportions of other immune cells (B-cells, DC, Monocytes/macrophages) were also noticed in this donor. This donor was the only one subjected to MACS dead cell removal, which was later excluded from the protocol due to concerns about losing larger cell types such as upper epithelial cells (0.5% of all cells in 296C, over 7% in all other donors). In addition, this donor was diagnosed with ventilator-associated pneumonia and some reports in mice indicate a link between mast cells and pneumonia infection (38,39).

All the 94,257 cells from spleen were annotated as immune cells. Follicular and mantle zone B-cells were identified as the largest group with 17% (>16,000 cells) and 20% (>18,000 cells), respectively. Dividing B-cells potentially undergoing affinity maturation, were annotated by the expression of AICDA and detected with a frequency of 0.5% (437 cells). Over 6000 plasma cells were detected and annotated as plasmablasts, IgG or IgM expressing plasma cells. About 90% from each of those originated from one donor 356C, which is consistent with the medical records showing chest infection in this donor. Over 28,000 T-cells were annotated as CD4+ conventional, CD8+ activated, CD+4 naive, CD4+ follicular helper (CD4_fh), CD8+ MAIT-like, CD8+ gamma-delta, CD8+ cytotoxic lymphocyte (CD8 CTL), CD4+ regulatory or dividing T-cells. Two subpopulations of natural killer (NK) cells, a dividing NK population, monocytes, macrophages and DC-s were also identified. Multiple cell groups were represented in very low proportions, such as sub-populations of the DC including activated DC (0.04%), conventional DC1 (0.3%) and pcDC-s (0.3%), as well as Innate Lymphoid Cells (0.6%), CD34+ progenitor cells (0.2%), platelets (0.08%) and an unknown population of cells positioned between T and B cells clusters (0.1%). Another group containing over 2207 cells expressing both T- and B-cells markers were called T-cells/B-cells, and could represent the doublets of interacting cells. Besides the plasma cells populations, multiple other cell types such as B-cell/T-cell, conventional DC1 and DC2, pcDC and macrophages were also represented in higher proportions in donor 356C than any other donor. No stromal cells were detected, which is likely to be due to the fact that for spleen no enzymatic digestion was employed to release cells.

#### Tissue processing signatures

We also performed bulk RNA-sequencing for each donor at each time point to assess gene expression changes over time without dissociation artefacts and to allow us to determine what gene sets are changed by dissociation, or if specific cell populations are lost. On a UMAP plot, the bulk and single-cell pseudo-bulk (sc-pseudo-bulk) samples cluster primarily by method (bulk or sc-pseudo-bulk) and by tissue of origin, but not according to the time point (Supplementary Figure 9). Previous work has highlighted the effect of enzymatic tissue dissociation on gene expression patterns (20). Differential expression analysis was carried out by Wilcoxon signed-rank test in each tissue between bulk vs sc-pseudo-bulk samples. None of the Bonferroni FDR corrected p-values from pairwise comparisons passed the significance threshold (adj. p-val < 0.05) in any of the tissues. However, the genes with the highest fold changes from sc-pseudo-bulk to bulk had significant p-values without multiple testing, and were enriched in ribosomal genes in all three tissues (Supplementary Table 5). Also, a long non-coding RNA, MALAT1, appeared in the top 20 genes expressed at higher levels in sc-pseudo-bulk in all of the three tissues. The high enrichment of ribosomal genes in sc-pseudo-bulk samples was also evident when combining all tissues for the analysis. Over 16,000 genes were significantly different after multiple testing correction with this increased sample size. The list of highly expressed genes in sc-pseudo-bulk was enriched with ribosomal genes (adjusted p-val 1.15 e-06), as well as MALAT1 (adjusted p-val 1.15 e-06, log2 fold change 4.4). Nearly all of the dissociation-related FOS, FOSB, JUN and JUNB genes (20) were also significantly higher in the sc-pseudo-bulk than in the bulk samples with adjusted p-values 0.002, 2.15 e-06, 0.15 (not adjusted p-value 4.49 e-06) and 7.65 e-05, respectively. Differential expression of ribosomal and early response genes was also seen in the dissociation signature reported by Van Oudenarden (20).

We also carried out tissue specific analysis of differential gene expression. Genes more highly expressed in bulk derive from the cell types sensitive to dissociation. Pulmonary alveolar cells are very scarce in our single-cell lung data, but abundant in the tissue. This results in the differential expression of the marker AGER and surfactant-protein encoding genes SFTPB, SFTPA1 and SFTPA2. Other genes with high fold changes between bulk and sc-pseudo-bulk are blood vessel endothelial markers VWF and PECAM1. In oesophagus, stromal specific genes FLNA and MYH11 and both KRT4 and KRT5, expressed in the majority of keratinocytes, are higher in bulk vs sc-pseudo-bulk. In spleen, the list of top genes includes APOE, CD5L, VCAM1, HMOX1, C1QA and C1QC, which are strongly expressed in macrophages. This suggests that our sample processing protocols are mostly affecting alveolar and blood vessel cells in lung, stromal cells in oesophagus and macrophages in spleen. However, none of these genes are significant after multiple testing correction. Importantly for this study, these genes were upregulated by the dissociation process, but this upregulation was not altered by cold storage time prior to dissociation.

#### Cell type specific changes in transcriptome

Having annotated cell types, it was possible to examine whether there was a cell type specific effect of storage time. Notably, UMAP plots did not reveal an obvious effect of time (Figure 4g-h). We joined gene expression matrices for all the tissues and calculated the percentage of variability explained by different variables. Figure 4j shows that donor, tissue, cell type and number of counts account for a large fraction of the variance while the effect of storage time made the smallest contribution. This remained the case when the analysis was carried out per tissue (Supplementary Figure 10).

We next examined whether the observed increase in mitochondrial reads with time (spleen, 72 hours, Figure 2e-g) was due to a specific cell type. For this purpose, cells with high mitochondrial reads were also assigned to a cell type via similarity. For each cell type and tissue, the mitochondrial percentages and their fold changes relative to T0 were calculated (Supplementary Figure 10, Figure 5). The highest fold changes and significance were present in spleen at 72h. While this effect was apparent in multiple cell types, it was particularly evident in plasma cells; being independently replicated in the two donors contributing the majority of this cell type. (Supplementary Figure 11, Supplementary Figure 12).

**Figure 5:**
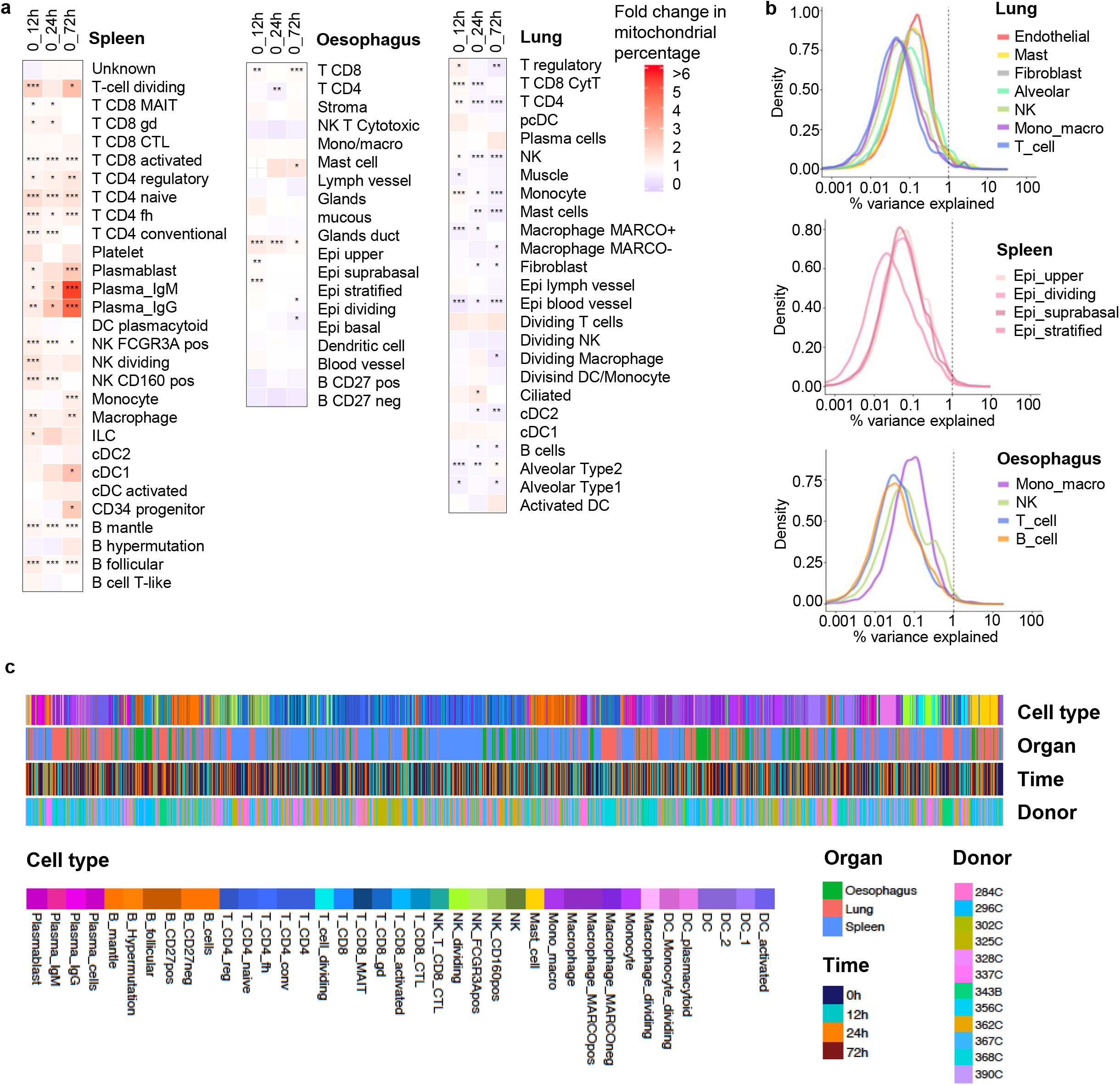
Cell type specific changes in transcriptome. (a) Proportion of mitochondrial reads relative to T0 calculated for spleen, oesophagus and lung. The fold change (FC) of mitochondrial percentage is measured in every cell type between T0 and 12h, 24h and 72h. FC is indicated by color with white indicating no fold change (FC=1), blue indicates a drop in mitochondrial percentage, red indicates an increase in mitochondrial percentage compared to T0 (FC>1). Benjamin and Hochberg adjusted p-values are indicated by asterisk as follows: p-val<0.01*, p-val < 0.00001** and p-val < 0.00000001***. All cells are used including the ones with high mitochondrial percentage (>10%), annotated via scmap tool. Grey indicated time points with fewer than 5 cells. Missing values (no sample) are shown by cross. (b) Percentage of variance in gene expression explained by time for cell type groups in lung, oesophagus and spleen. Cell type groups in lung are Endothelial (blood vessel, lymph vessel), Alveolar (alveolar Type 1 and Type 2), Mono_macro (Monocyte, Macrophage_MARCOneg, Macrophage_MARCOpos), T_cell (T_CD4, T_CD8_Cyt, T_regulatory). Cell type groups in spleen are Mono_macro (Monocyte, Macrophage), NK (NK_FCGR3Apos, NK_CD160pos), T_cell (T_CD4_conv, T_CD4_fh, T_CD4_naive, T_CD4_reg, T_CD8_activated, T_CD8_CTL, T_CD8_gd, T_CD8_MAIT-like, T_cell_dividing) and B_cell (B_follicular, B_Hypermutation, B_mantle). (c) Hierarchical clustering of cell types of up to 10 cells per cell type per tissue per donor and time. Cell attributes (cell type, organ, time and donor ID) are indicated by colour.

Next, similar cell types were joined together into larger clusters for more reliable analysis. The percentage of variability explained by time point in each of these cell type clusters was extremely low (Figure 5b), especially compared with variables such as donor and n counts (Figure 4j), highlighting that for almost all cell types cold storage time did not have a major effect.

We also examined which genes changed most with time in each cell type (see methods). Signatures were more similar within a tissue rather than within cell type, such as T-cells, natural killer cells and monocytes/macrophages from different organs (Supplementary Figure 13). Furthermore, the genes driving this similarity were among the top genes contributing to the ambient RNA contamination in the majority of samples (Supplementary Figure 14, Supplementary Table 6). Therefore, low level gene expression changes that we do observe with time are likely to be driven by cell death leading to ambient RNA contamination.

Pairwise differential expression analysis in bulk RNA-sequencing between T0 and other time points did not yield significant genes in any tissue (Supplementary Table 6), further indicating that any changes observed are extremely small. For lung we were also able to freeze samples at the clinic immediately after collection, and compare this sample to later time points. Again no significantly differentially expressed genes were detected.

It may seem surprising to observe so few changes in gene expression with time, especially given that other studies such as the GTEx project do demonstrate such effects (21,31). However it is important to note that post-mortem samples from warm autopsy were used for the Genotype-Tissue Expression (*GTEx*) project (albeit with <24h PMI). Our study was designed to mimic the process used during organ transplantation, in which tissues are removed rapidly (within one hour of cessation of circulation) from cold-perfused donors and stored at 4°C in hypothermic preservation media such as University of Wisconsin (timing is tissue-dependent; heart can be stored for 4-6h, lungs median 6.5h, kidneys median 13h). Indeed, for some organs, there is evidence that they remain functional for longer (28,29). Further, the work of Wang et al (30), which looked at hypothermic preservation of mouse kidneys in HypoThermasol FRS, also demonstrated little change in gene expression over 72h. Therefore, while it is certainly true that rapid gene expression changes will occur under certain storage conditions, at least for the organs tested in this study, it appears these can be limited by maintaining the samples cold in hypothermic preservation media. Altogether, this will be very useful for designing further studies with fresh biological samples (including biopsies from living donors) with regards to sample collection time in clinic, transport to the lab, and storage until processing is convenient.

#### Mapping of cell types across organs

Having generated data sets for oesophagus, lung and spleen, we examined if cell types that can be found in all three organs, such as immune cells, would cluster by organ or by cell type. Figure 5c shows the result of hierarchical clustering using the 1000 most highly variable genes in up to 10 cells per cell type, tissue, time and donor. In this analysis of approximately 7500 cells, we see clear subclusters of mast cells, macrophages and plasma cells with some substructure depending on the donor and the tissue of origin, suggesting that more extensive analysis will allow us to study tissue adaptation of different immune cell populations. Other cells such a B cells sit in two groups, and dividing cells (NK, macrophages, T cells) also co-segregate. Importantly for this study, samples do not cluster by time.

#### Variation in cell type contribution

Our protocols of single cell dissociation are aimed at capturing the diversity of cell types present in each organ, but do not represent the proportion of each cell type in the original tissue. For example, the tissue dissociation protocol employed for lung strongly enriches for immune cell types. Relatively high variability in the proportions of cell types was seen between samples. This was likely to be due to some technical variation as well as underlying biological variation as indicated by the capture of rare structures such as glandular cells in only some oesophagus samples, namely from donor 325C and 356C. Interestingly, histology on sections from donor 325C (12h) confirmed the presence of glands (mucous secreting cells; Supplementary Figure 1) that were not present in the other oesophagus samples sectioned. All other samples contained fewer than 5 mucous secreting cells (Supplementary Table 2). This exemplifies the difficulty in collecting cells from structures that are sparsely distributed, such as the glands in the oesophagus, and suggests that some of the sample to sample variation is due to the underlying differences in the architecture of the specific tissue sections analysed. A similar effect was seen for lymph vessels (Figure 4e, 367C, 72h). Furthermore the immune infiltrate seen on one of the histology sections (Supplementary Figure 1c, 362C, 24h) is possibly reflected in an increase in immune cells (B, T and monocytes/macrophages) at the single cell level (Figure 4d, 362C, 24h).

Overall, we observe greater variability between donors than between samples and the prevalence of certain cell types in multiple samples from the same donor supports this conclusion. For example, all lung samples from donor 356C were highly enriched for NK cells, while those from donor 368C contained high numbers of macrophages (Figure 4 d-f). This high variability between donors suggests that for the Human Cell Atlas a large number of donors will have to be profiled to understand the range of “normal”.

## Conclusions

We present a method for cold storage of primary human tissue samples that requires no processing at the clinical site beyond sample collection, and permits at least a 24h window for shipping, tissue dissociation and scRNA-seq. Lung and oesophagus appeared stable for 72h by all of the metrics tested. In spleen we observe changes in the proportion of intronic and exonic reads and an increase in the percentage of mitochondrial reads at 72h. We see no effect of time on the diversity of cell populations in scRNA-seq data, or change in bulk RNA-seq. This method is easy to adopt and will greatly facilitate primary sample collections for Human Cell Atlas studies. We highlight changes of cell type distribution due to anatomical heterogeneity of tissue samples and significant heterogeneity between donors, which will impact future HCA study design.

Further, we have generated detailed annotations on three primary human tissues: spleen, oesophagus and lung. This dataset of over 240,000 single cells presents a significant resource for further investigation of the biology of these tissues and contains the largest oesophagus and spleen datasets to date. In addition, we make available WGS data from thirteen healthy donors including clinical metadata, allowing for future tissue-specific, single cell eQTL studies.

## Methods

### Aim and study design

We aimed to identify a method of preserving intact human tissue samples for scRNAseq. Three tissues, expected to have different sensitivities to ischaemia (n=5-6 per tissue), were chosen: spleen, oesophagus and lung. One sample was processed for 10x Genomics 3’ scRNAseq (v2) immediately upon receipt (T0), and the remainder processed after 12h, 24h and 72h cold ischaemic time. Additional samples were collected for bulk RNA extraction at each time point, and genomic DNA also prepared for WGS from each donor. Of note, an additional lung donor was collected (376C) which is not analysed but is available in the Data Co-ordination Platform submission. Upon receipt, this sample was morphologically abnormal (blackened) and the resulting cell suspension contained cells with black granules, likely due to the donor being a heavy smoker for a prolonged period.

### Donor samples

All samples were obtained from the Cambridge Biorepository for Translational Medicine (CBTM) under appropriate ethical approval (see ‘Ethics approval and consent to participate’). These were primarily from donation after circulatory death (DCD) organ donors, in whom circulatory arrest followed withdrawal of life-sustaining treatment. Patient characteristics are listed in Table 4 and representative histology is shown in Supplementary Figure 1. Upon cessation of circulation, donors were perfused with cold University of Wisconsin (UW) solution and research samples collected at the end of the transplant procedure, within 1-2 hours of cessation of circulation, constantly under cold ischaemic conditions. Samples (typically 0.5-2cm^3^) were maintained on ice in UW in the operating theatre, then rapidly transferred into cold HypoThermasol FRS preservation solution at 4°C for shipping / storage (Sigma H4416). On receipt at the processing laboratory (typically 4h following cessation of circulation) these were dissociated for 10x Genomics 3’ single cell sequencing (v2), and a portion flash frozen in isopentane for bulk RNA / DNA extraction as soon as possible (‘T0’ time point), or at 12, 24 and 72h ischaemic time following storage in the fridge (4°C). For lung samples it was also possible to collect an additional flash frozen sample at the clinic immediately after tissue excision, in order to compare bulk RNA between this ‘true zero’ time and the ‘T0’ time point, the latter being frozen on receipt at the tissue processing laboratory. The start of ischaemia was defined as the point at which circulation ceased for donation after cardiac death (DCD) donors, unless they received normothermic regional perfusion with oxygenated blood (NRP, 2h), in which case the end of NRP was used. For the one donation after brain stem death (DBD) donor in the study, start of ischaemia was defined as the time at which life support was withdrawn. End of ischaemia was defined as time of cell lysis or freezing; for 10x single cell reactions lysis occurs in the PCR step immediately after loading on 10x Genomics Chromium instrument. Cold ischaemic times are available in the Data Co-ordination Platform metadata submission.

### Tissue dissociation

All tissue dissociation protocols are available on protocols.io (40): spleen (protocol 32rgqd6), oesophagus epithelium (protocol 34fgqtn) and lung parenchyma (protocol 34kgquw).

Spleen (protocols.io 32rgqd6): samples from the top 5-7mm of the organ were mechanically mashed through a 100μM cell strainer with cold PBS, pelleted at 500xg, 5 min and resuspended in cold 1x red blood cell lysis buffer (Life Technologies). Following dilution in cold PBS, pelleting at 500xg, 5 min and resuspension in cold 0.04% BSA / PBS, cells were counted and viability was determined using a C-chip haemocytometer and trypan blue. Up to 10 million cells were used for MACS dead cell removal (Miltenyi; protocols.io qz5dx86) and the flow through (live cells) pelleted, resuspended in cold 0.04% BSA / PBS and counted / viability determined using trypan blue and C-chip. Cells were loaded onto the 10x Genomics Chromium Controller following the single cell 3’ v2 protocol, aiming for between 2000-5000 cells recovery.

Oesophagus epithelium (protocols.io 34fgqtn): a cylindrical piece of the oesophagus was received from the mid-region, the mucosa (mainly epithelium) removed mechanically with forceps / scissors and divided into segments for time points (placed in HypoThermasol FRS in the fridge). The epithelium / mucosa was finely chopped with scalpels and incubated for 30min in 0.25% trypsin-EDTA (GIBCO) containing 100μg/ml DNase I (Sigma) at 37°C with shaking. The sample was centrifuged and digestion media replaced with fresh 0.25% trypsin-EDTA (GIBCO) / DNase I for 15min at 37°C with shaking (this protocol can also be used for stomach, in which the media change is necessary due to pH alterations as the tissue digests; this is less required for oesophagus). Trypsin was neutralised with RPMI containing 20% FBS, cells pelleted and passed through a 70μM strainer before pelleting again and treating with 1x red blood cell lysis buffer (Life Technologies). Following dilution, pelleting and resuspension in cold 0.04% BSA / PBS, cells were counted and viability determined using a C-chip haemocytometer and trypan blue. The resulting suspension contained a range of cell sizes, up to 50μM. No dead cell removal was performed for oesophagus samples due to the risk of losing larger cells in the MACS column (cell viability was >70%), with the exception of the fresh sample from the first oesophagus donor (296C). Cells were loaded onto the 10x Genomics Chromium Controller following the single cell 3’ v2 protocol, aiming for 5000 cells recovery.

Lung (protocols.io 34kgquw): a 0.2-0.5g piece of lung parenchyma (lower left lobe) was finely chopped with scalpels and incubated for 1h in 0.1mg/ml collagenase D (Sigma C5138) in DMEM with 100μg/ml DNase I (Sigma) for 1h at 37°C with shaking. (This protocol was initially designed for isolation of immune cells from lung airway, a much tougher region of lung tissue; parenchyma can be dissociated with 30 min treatment; however 1h incubation was used for this study). Digested tissue was mashed through a 100μM cell strainer and washed with DMEM containing 10% FBS before centrifuging, washing and resuspending the pellet in 1x red blood cell lysis buffer (Life Technologies). Following dilution, pelleting and resuspension in cold 0.04% BSA / PBS, cells were counted and viability determined using a C-chip haemocytometer and trypan blue. MACS dead cell removal was performed (Miltenyi; protocols.io qz5dx86) and the flow through (live cells) pelleted, resuspended in 0.04% BSA / PBS and counted using trypan blue and C-chip. Cells were loaded onto the 10x Genomics Chromium Controller following the single cell 3’ v2 protocol, aiming for 5000 cells recovery.

### Library preparation, bulk RNA and WGS

cDNA libraries were prepared from single cell suspensions following the 10x Genomics 3’ v2 protocol and 2 samples per lane sequenced on HiSeq4000 with 26bp read 1, 8bp sample index and 98bp read 2 (aiming for 150M reads / sample or ≥30,000 per cell).

For bulk RNA and DNA extraction, samples were flash frozen in isopentane at each time point (protocols.io qz7dx9n). Bulk RNA and DNA was prepared from frozen samples using the Qiagen AllPrep DNA/RNA mini kit and TissueLyser II. Spleen RNA samples required an additional on-column DNase digest.

RNA was quantified using the QuantiFluor RNA system (Promega) on a Mosquito LV liquid handling platform, Bravo WS and BMG FluoSTAR Omega plate reader. Libraries (poly(A) pulldown) were prepared using the NEB RNA Ultra II Custom kit on an Agilent Bravo WS automation system, including PCR with the KAPA HiFi Hot Start Mix and dual-indexing. Libraries were cleaned on a Caliper Zephyr liquid handling system using Agencourt AMPure XP SPRI beads and quantified with the AccuClear™ Ultra High Sensitivity dsDNA Quantitative kit (Biotium). RNA integrity number (RIN) was determined for each sample by Agilent BioAnalyser RNA 6000 Nano kit. Libraries were pooled in equimolar amounts and quantified on an Agilent BioAnalyser before sequencing on an Illumina HiSeq4000, 75bp paired end, aiming for 35 million reads per sample.

Genomic DNA from 13 donors was prepared for WGS. DNA was first sheared to 450bp using a Covaris LE220 instrument, purified with AMPure XP SPRI beads (Agencourt) on an Agilent Bravo WS and then libraries prepared with the NEB Ultra II custom kit on an Agilent Bravo WS system. PCR (6 cycles) was performed using the Kapa HiFi Hot Start Mix and IDT 96 iPCR tag barcodes, before purification using Agencourt AMPure XP SPRI beads on a Beckman BioMek NX96 liquid handling platform. Libraries were sequenced at 30x coverage on an Illumina HiSeqX.

### Computational analysis

#### Single cell RNA-seq data analysis

Reads were mapped to GRCh38 1.2.0 Human Genome reference by Cellranger 2.0.2 pipeline. The EmptyDrops algorithm (41) was run on each sample. Identified cells were used to generate the Cellranger filtered count matrix. An outlier sample HCATisStabAug177276393 (spleen, Donor 302C, 24h) in which fewer than 40% of reads mapped to the transcriptome was removed from further analysis (Supplementary Figure 2a). Count matrices were analysed by scanpy version 1.4 (42) tool in Python version 3.7.2. Cells with less than 300 or more than 5000 detected genes (8000 in oesophagus), more than 20,000 UMI and more than 10% mitochondrial reads were removed. Genes that were detected in less than three cells per tissue were removed. All donors and time points per tissue were combined for analysis. The reads were log-transformed and normalised.

#### Quality metrics of samples

Number of cells, number of reads, median genes per cell, reads confidently mapped to the transcriptome and other quality metrics were obtained from Cellranger’s output metrics.csv files. The confidently mapped reads to intronic, exonic and intergenic regions were further studied by extracting the number of reads mapping confidently (QC=225 from Cellranger’s output bam file) for every cell barcode.

#### UMI counts analysis

Number of UMIs in each droplet was quantified by using soupX tool (37) in R. Debris was defined as reads in droplets that contained between 3 and 100 UMIs. The droplets containing 1 or 2 UMIs were defined as ambient RNA expression originating from free-floating RNA in the sample. Droplets containing over 100 counts were defined as cellular material.

#### Clustering and annotation of cell types

To achieve good clustering by cell types, number of counts, mitochondrial percentage and donor effects were regressed out. PCA was carried out on highly variable genes, the donor effect was reduced by BBKNN tool (43). Leiden clustering (44) and UMAP visualisation was performed for gaining clusters of cells and visualisation. Statistical analysis was performed in R version 3.5.0 and plotting was in Python via scanpy or custom script and in R using ggplot2 version 2.2.1 or by using custom scripts. Cells which contained more than 10% mitochondrial reads were assigned by similarity to their closest cell type within a tissue with scmap tool (45), using cells with less than 10% mitochondrial reads as a reference. The high and low mitochondrial percentage cells were then combined for calculating the mitochondrial percentage per each cell type. All code for the analysis is available at https://github.com/elo073/TissStab.

Expression of known markers and re-analysis of bigger clusters were used to annotate cell types, with cell markers shown in Supplementary Figure 7. The major cell types were annotated for lung, oesophagus and spleen by looking at expression of known cell type markers. Three subsets from lung (mononuclear phagocytes and plasma cells; lymphocytes; dividing cells), two subsets from oesophagus (Immune; small clusters) and two subsets from spleen (DC, small clusters and dividing cells; CD4 and CD8 T-cells) were extracted, further analysed by re-clustering and annotated using known markers. These updated annotations then replaced the original bigger ones.

#### Explanatory variance calculation

Effect of variable factors (donor, tissue, time point, cell type, n_counts, etc) on gene expression was studied by the scater package by computing the marginal R^2^ that describes the proportion of variance explained by each factor alone for each gene. Density plots of the gene-wise marginal R^2^ are shown. Normalised and scaled gene expression was used with the effect of donor and number of counts regressed out, but not mitochondrial percentage or time. Effect of time only as a continuous variable was calculated for each cell type or cell type group in tissues. Smaller or similar cell types were combined to groups as Endothelial (blood vessel, lymph vessel), Alveolar (alveolar Type 1 and Type 2), Mono_macro (Monocyte, Macrophage_MARCOneg, Macrophage_MARCOpos), T_cell (T_CD4, T_CD8_Cyt, T_regulatory) in Lung and Mono_macro (Monocyte, Macrophage), NK (NK_FCGR3Apos, NK_CD160pos), T_cell (T_CD4_conv, T_CD4_fh, T_CD4_naive, T_CD4_reg, T_CD8_activated, T_CD8_CTL, T_CD8_gd, T_CD8_MAIT-like, T_cell_dividing) and B_cell (B_follicular, B_Hypermutation, B_mantle) in Spleen.

#### Differential expression

Wilcoxon signed-rank test was used to compare expression between time points in bulk RNA-sequencing samples, and between bulk RNA-sequencing and sc-pseudo-bulk samples. Bonferroni corrected FDR values were reported, as well as the median_log2_foldchange.

## Abbreviations

eQTL: Expression quantitative trait loci
scRNAseq: Single cell RNA sequencing
HCA: Human Cell Atlas
DMSO: Dimethyl sulfoxide
GTEx: Genotype-Tissue Expression
CBTM: Cambridge Biorepository for Translational Medicine
DCD: Donation after cardiac death
DBD: Donation after brainstem death
UW: University of Wisconsin
NRP: Normothermic regional perfusion
PCR: Polymerase chain reaction
RNA: Ribonucleic acid
DNA: Deoxyribonucleic acid
PBS: Phosphate buffered saline
BSA: Bovine serum albumin
MACS: Magnetic activated cell sorting
FBS: Foetal bovine serum
HTA: Human Tissue Authority
REC: Research Ethics Committee
PMI: Post mortem interval
WGS: Whole Genome Sequencing
Sc-pseudo-bulk: Single-cell pseudo-bulk

## Declarations

### Ethics approval and consent to participate

Tissue samples were obtained from the Cambridge Biorepository for Translational Medicine (CBTM; (46)) with informed consent from the donor families and approval from the NRES Committee of East of England – Cambridge South (15/EE/0152). This consent includes generation of open-access genetic sequencing data and publication in open access journals in line with Wellcome Trust policy. CBTM operates in accordance with UK Human Tissue Authority guidelines (HTA; 47).

### Consent for publication

All samples were provided as linked-anonymised, with the linkage key held by the clinician. Consent was obtained as part of the Research Ethics Committee (REC) approval.

### Availability of data and material

The datasets generated in this study are available through the Human Cell Atlas Data Co-ordination Platform: (https://prod.data.humancellatlas.org/explore/projects/c4077b3c-5c98-4d26-a614-246d12c2e5d7). The data can also be browsed at: www.tissuestabilitycellatlas.org. All experimental protocols are available on Protocols.io, https://www.protocols.io/ (40).

### Authors’ contributions

EM performed the bulk of the data analysis. AWC, MJTS and OS designed the study. AWC, RJM, PH, KL, and AT generated the experimental data. KSP, KM and NG collected the samples. MD performed histology. RJM, KP, RA and SVD contributed to data analysis. EM, AWC and KBM wrote the manuscript. KNO, RF, JF, MC and MP assisted with tissue dissociation protocols, laboratory set up and provided substantial advice throughout the data generation and analysis process. ST and KBM provided critical evaluation of the data and manuscript. All authors read and approved the final manuscript.

## Acknowledgements

We would like to thank the donors and their families for the generous gift of tissue to this research project and members of the Cambridge Biorepository for Translational Medicine, who collected the samples. Also, the Cellular Genetics and Phenotyping (CGAP) core facility at the Sanger Institute, especially Adam Hunter, Liam Bolt, Emily Relton and Cecilia Icoresi Mazzeo for help with tissue processing. Lia Campos helped with assessing the histological stainings. Vladimir Kiselev and Anton Kodak at Sanger Institute Cellular Genetics Informatics team, Ricard Argelaguet from the European Bioinformatics Institute and Jongeun Park from the Sanger Institute supported the analysis with computing environment and scripts. Finally, the Sanger Institute DNA Pipelines facility prepared and sequenced the libraries.

