## Supplementary Figures for "Lung, spleen and oesophagus tissue remains stable for scRNAseq in cold preservation"

### Table of contents

**Title**: Lung, spleen and oesophagus tissue remains stable for scRNAseq in cold preservation 1

**Table of contents** **2**

**Supplementary Table legends** **3**

Supplementary Table 1. Patient characteristics. 3

Supplementary Table 2. Number of cells in cell types. 3

Supplementary Table 3. RIN-values for samples. 3

Supplementary Table 4. Differential Expression between the bulk and single-cell pseudo-bulk RNA-sequencing samples. 3

Supplementary Table 5. Pairwise Differential Expression between time points with bulk RNA-sequencing data in three tissues. 3

Supplementary Table 6. Explained variability by time in different cell types. 4

**Supplementary Figures** **5**

Supplementary Figure 1. Histological analysis. 5

Supplementary Figure 2. scRNA-seq quality metrics for all samples. 7

Supplementary Figure 3. Fraction of reads in exonic versus intronic regions changes in time in spleen, oesophagus and lung. 8

Supplementary Figure 4. Doublet score predictions do not change with storage time. 9

Supplementary Figure 5. Proportions of droplets in three intervals. 10

Supplementary Figure 6. Cell viability directly after dissociation and after dead cell removal. 11

Supplementary Figure 7. Cell type markers and their expression in the data. 12

Supplementary Figure 8: Distribution of cells from different donors. 15

Supplementary Figure 9. Bulk RNA-sequencing data comparison with single-cell RNA-sequencing. 16

Supplementary Figure 10. Time explains the least of variance in gene expression. 17

Supplementary Figure 11. Mitochondrial percentage differs between cell types. 18

Supplementary Figure 12. Change in the percentage of mitochondrial reads in time and by cell type and donor in spleen. 19

Supplementary Figure 13. Gene signatures associated with storage time are correlated with tissue type and not cell type. 20

Supplementary Figure 14. Ambient RNA contamination genes. 21

### Supplementary Table legends

##### Supplementary Table 1. Patient characteristics.

Selected patient metadata. DCD = Donation after cardiac death, DBD = Donation after brainstem death, NRP = normothermic regional perfusion. Additional metadata is available in the Data Coordination Platform submission, or upon request.

##### Supplementary Table 2. Number of cells in cell types.

Number of cells is given for each cell type in every sample according to the tissues. Samples are shown in columns as a combination of time and donor (time_donor), cell types are shown in rows. Tissues are distributed in three separate sheets of the workbook: Lung, Oesophagus and Spleen.

##### Supplementary Table 3. RIN-values for samples.

Quality of the samples was assessed by the RNA Integrity number (RIN) measured by Agilent Bioanalyser. RIN values are shown and plotted for all the samples that were analysed.

##### Supplementary Table 4. Differential Expression between the bulk and single-cell pseudo-bulk RNA-sequencing samples.

Single-cell pseudo-bulk (sc-pseudo-bulk) samples were compared with their corresponding bulk RNA-sequencing data. The pairs were subjected to the Wilcoxon signed-rank test. P-values and Bonferroni corrected FDR p-values are shown in columns “pvals” and “pvals_adj” correspondingly. Median log2 fold-change is shown for every gene in column “median_log2_foldchange”. Positive values imply upregulation in the first group listed, negative values imply upregulation in the second group tested. Four sheets in the workbook have results for four different comparisons: Lung bulk vs sc-pseudo-bulk, Oesophagus bulk vs sc-pseudo-bulk, Spleen bulk vs sc-pseudo-bulk and all tissues combined bulk vs sc-pseudo-bulk.

##### Supplementary Table 5. Pairwise Differential Expression between time points with bulk RNA-sequencing data in three tissues.

Wilcoxon signed-rank test was performed between the Clinic time point (“true 0h”) and all other time points (T0, 12h, 24h, 72h) in lung, and between T0 and all other time points (12h, 24h, 72h) in lung, oesophagus and spleen shown in four different workbook sheets. P-values and Bonferroni corrected FDR p-values are shown in columns “pvals” and “pvals_adj” correspondingly. Median log2 fold-change is shown for every gene in column “median_log2_foldchange”. Positive values imply upregulation in the first group listed, negative values imply upregulation in the second group tested.

**Supplementary Table 6. Explained variability by time in different cell types.**

Proportion of variance explained by time was calculated for each cell type or cell type group and is shown for genes that explained >= 1% of the variance in time in any of the cell types or cell type groups. Cell types were grouped as follows: Endothelial (blood vessel, lymph vessel), Alveolar (alveolar Type 1 and Type 2), Mono_macro (Monocyte, Macrophage_MARCOneg, Macrophage_MARCOpos), T_cell (T_CD4, T_CD8_Cyt, T_regulatory) in Lung and Mono_macro (Monocyte, Macrophage), NK (NK_FCGR3Apos, NK_CD160pos), T_cell (T_CD4_conv, T_CD4_fh, T_CD4_naive, T_CD4_reg, T_CD8_activated, T_CD8_CTL, T_CD8_gd, T_CD8_MAIT-like, T_cell_dividing) and B_cell (B_follicular, B_Hypermutation, B_mantle) in Spleen.

### Supplementary Figures

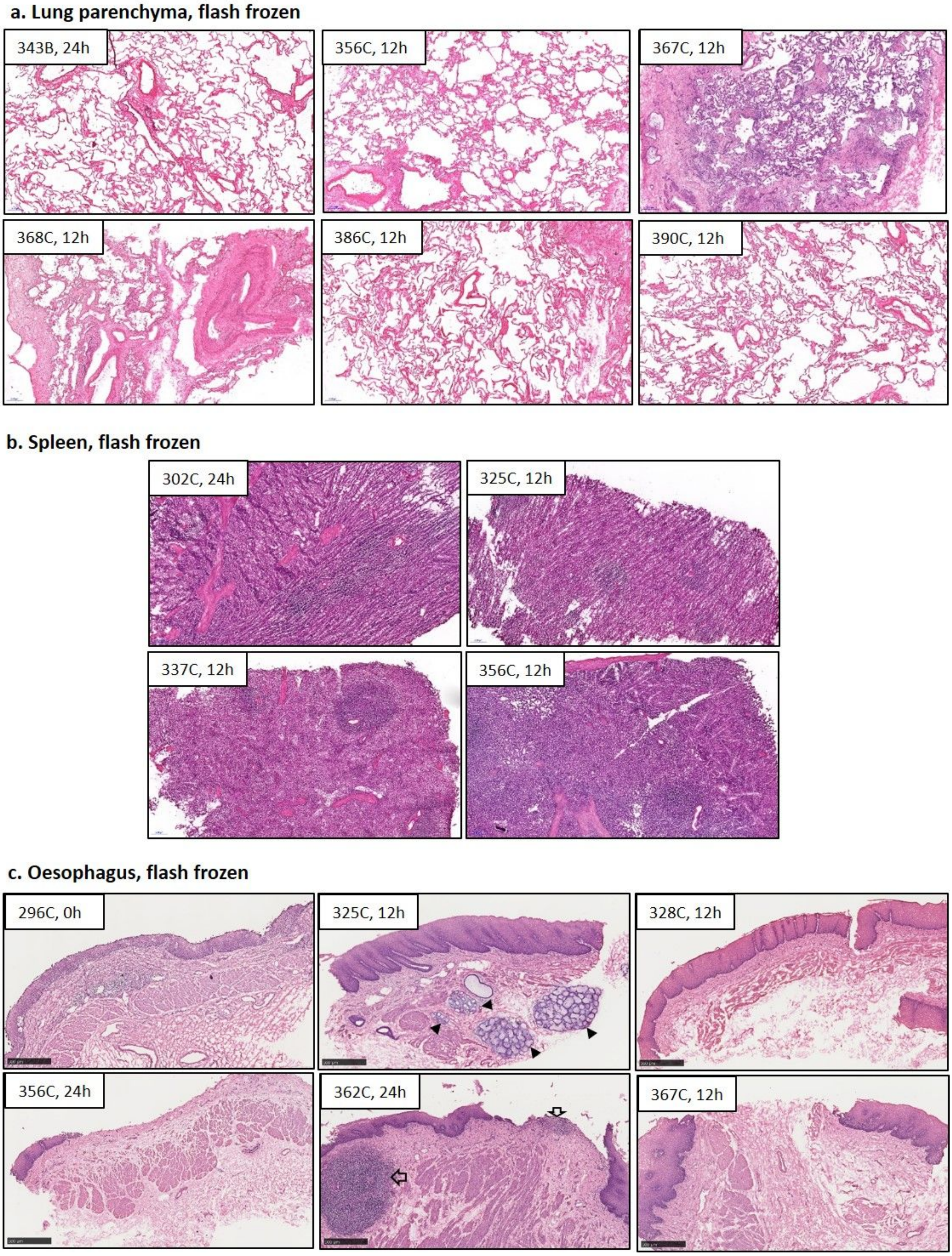


**Supplementary Figure 1. Histological analysis.**

Flash-frozen tissue samples from the time points indicated were cryosectioned and stained with haematoxylin and eosin for (**a**) lung, (**b**) spleen and (**c**) oesophagus. This shows normal, healthy histology overall (some emphysema is apparent in lung samples, but this is frequently observed in healthy patients). Of note, the lung from donor 368C exhibits some evidence of lung hypertension. A large vessel is apparent in the section taken from donor 368C. Note that apart from donor 296C, for which a full cross-section was retained, oesophagus samples contain only the mucosa / epithelium (the same layer used for single cell / bulk sequencing). Though damaged during processing in some samples, epithelium is clearly visible as a dark purple layer along the outer edge of the tissue. Oesophagus donor 325C, 12h, contains glands (arrow heads) and donor 362C, 24h has evidence of inflammation of unknown origin (open arrows). All images at 5x magnification; scale bar = 200µM for lung and spleen, 500µM for oesophagus.

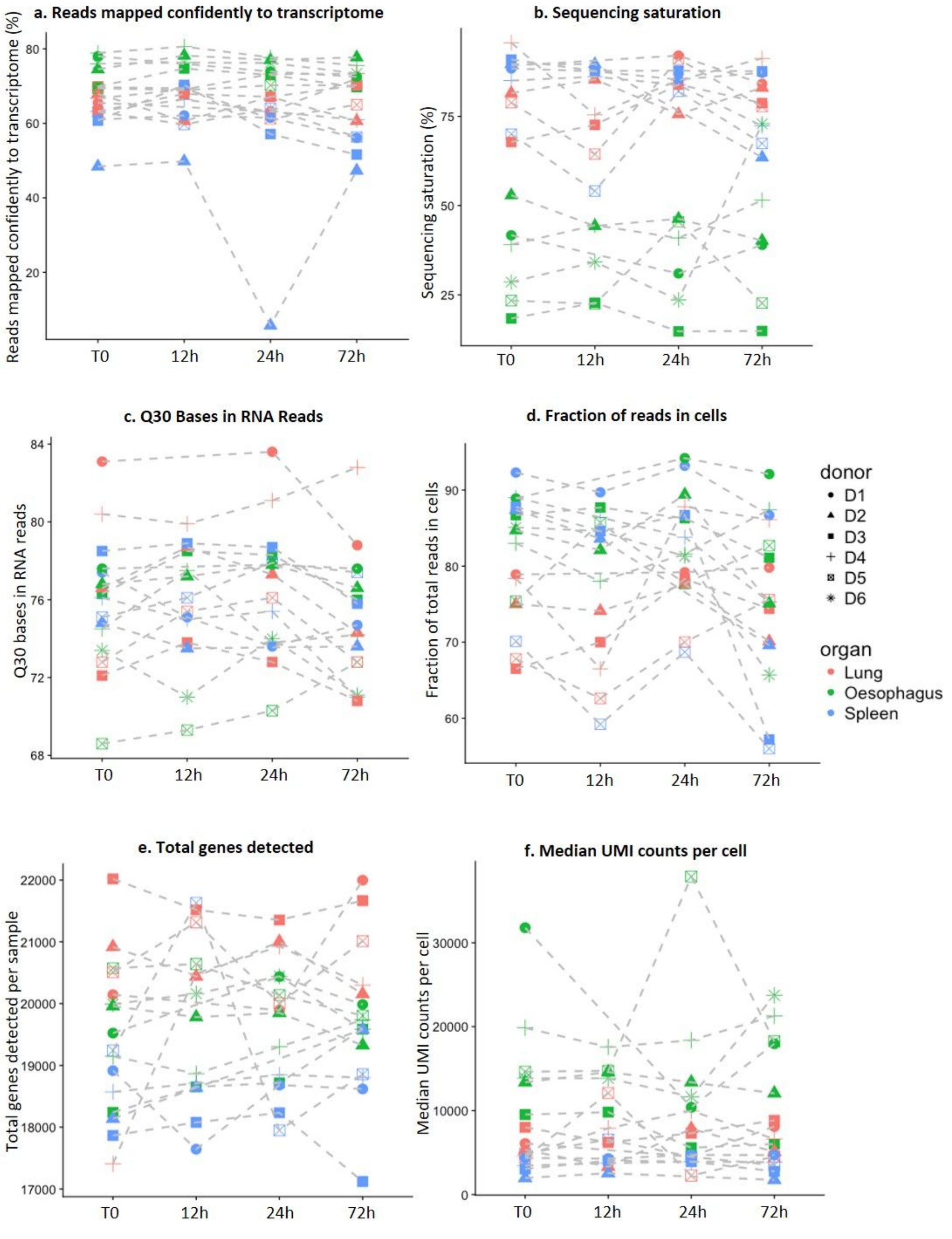


**Supplementary Figure 2. scRNA-seq quality metrics for all samples.**

Percentage of reads mapping confidently to the transcriptome (QC=225 from Cellranger 2.0.2 pipeline mapping the reads to GRCh38 1.2.0 Human Genome reference) shows an outlier with less than 40% (donor 2 for lung at time point 24h) and was removed from analysis (a). Percent sequencing saturation (b), Q30 bases in RNA reads (c), fraction of total reads assigned to cells (d), total number of genes detected per sample (e), median number of UMI counts per cell (f). Samples are colored by tissue, shapes correspond to a separate donor within the tissue.

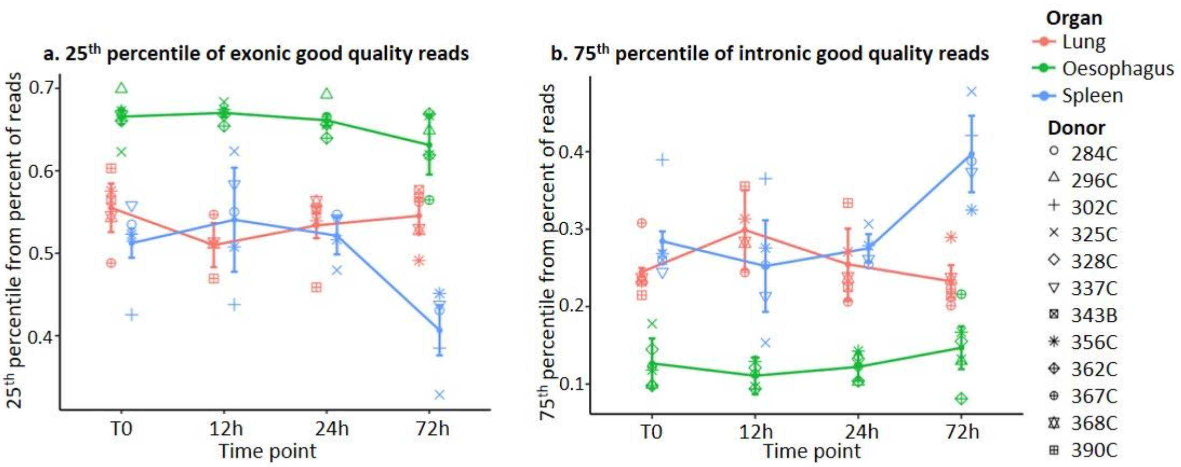


**Supplementary Figure 3. Fraction of reads in exonic versus intronic regions changes in time in spleen, oesophagus and lung.**

Percentage of good quality reads mapping to exons (a) or introns (b) in 25% or 75% of cells correspondingly. p-value is significant between T0 and 72h only in spleen for both exonic (p-value = 0.007) and intronic (p-value = 0.013) mapping fractions.

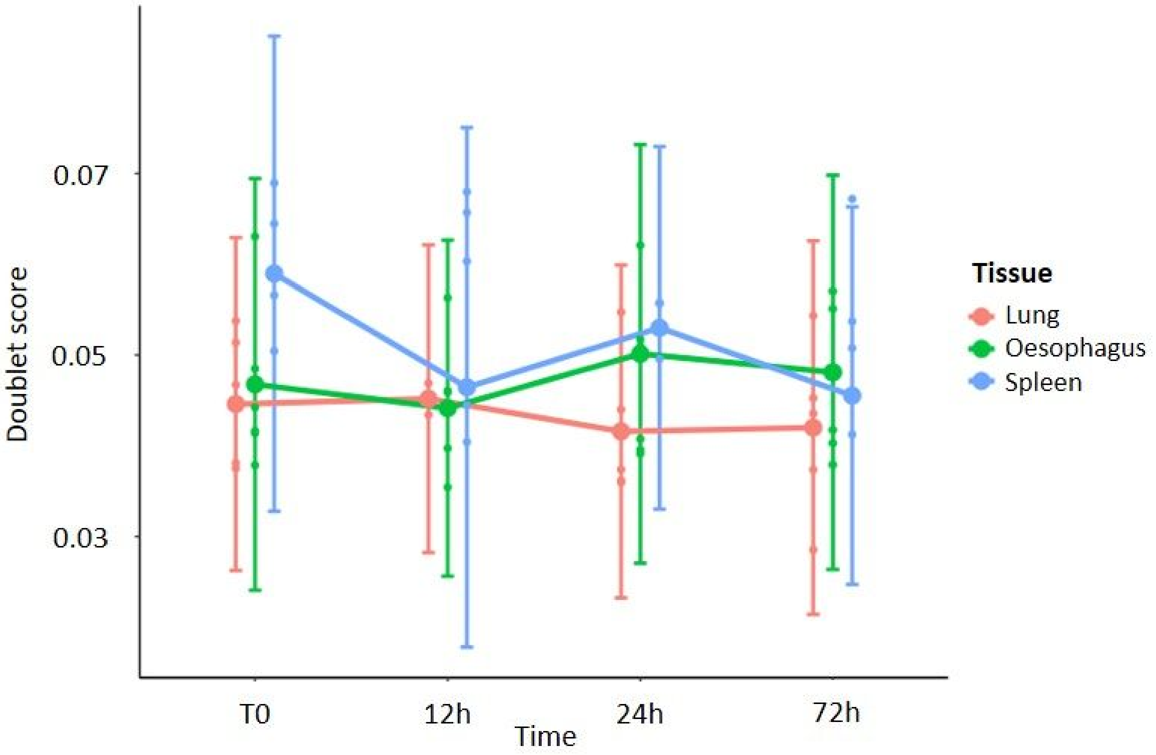


**Supplementary Figure 4. Doublet score predictions do not change with storage time.**

Scrublet was applied per every sample separately, doublets were simulated and soublet scores calculated. Mean values of doublet scores per run are plotted for each time point and tissue. Different tissues are shown by different colors, standard deviation is indicated by whiskers. Highest fold change is observed between T0 and 72h in spleen, providing p-value of 0.24.

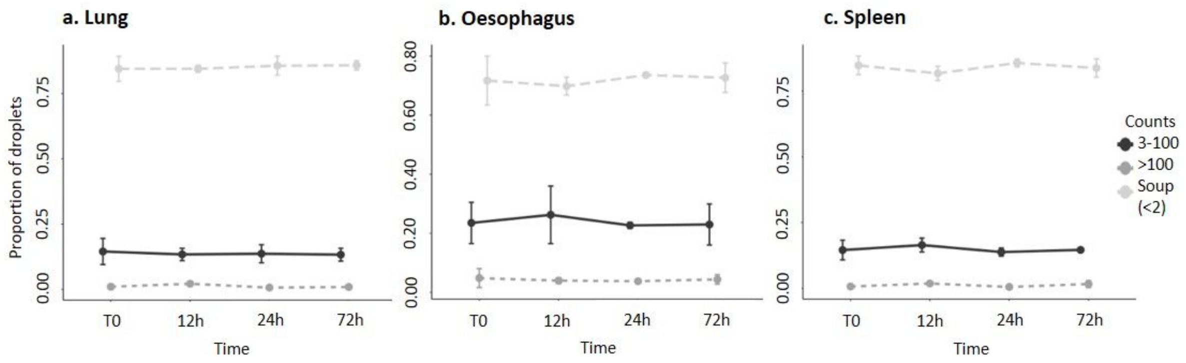


**Supplementary Figure 5. Proportions of droplets in three intervals.**

Proportion of droplets containing number of reads in given interval of UMI-s in Lung (a), Oesophagus (b) and Spleen (c) samples. Mean values and standard deviations are indicated by errorbar for donors within each tissue and timepoint, line types connect means within the same interval of UMI. Intervals of droplets containing 0 to 2 UMI counts (prop.soup), 3 to 100 counts (prop3_100) and more than 100 counts (prop100_higher) are shown by scale of grey.

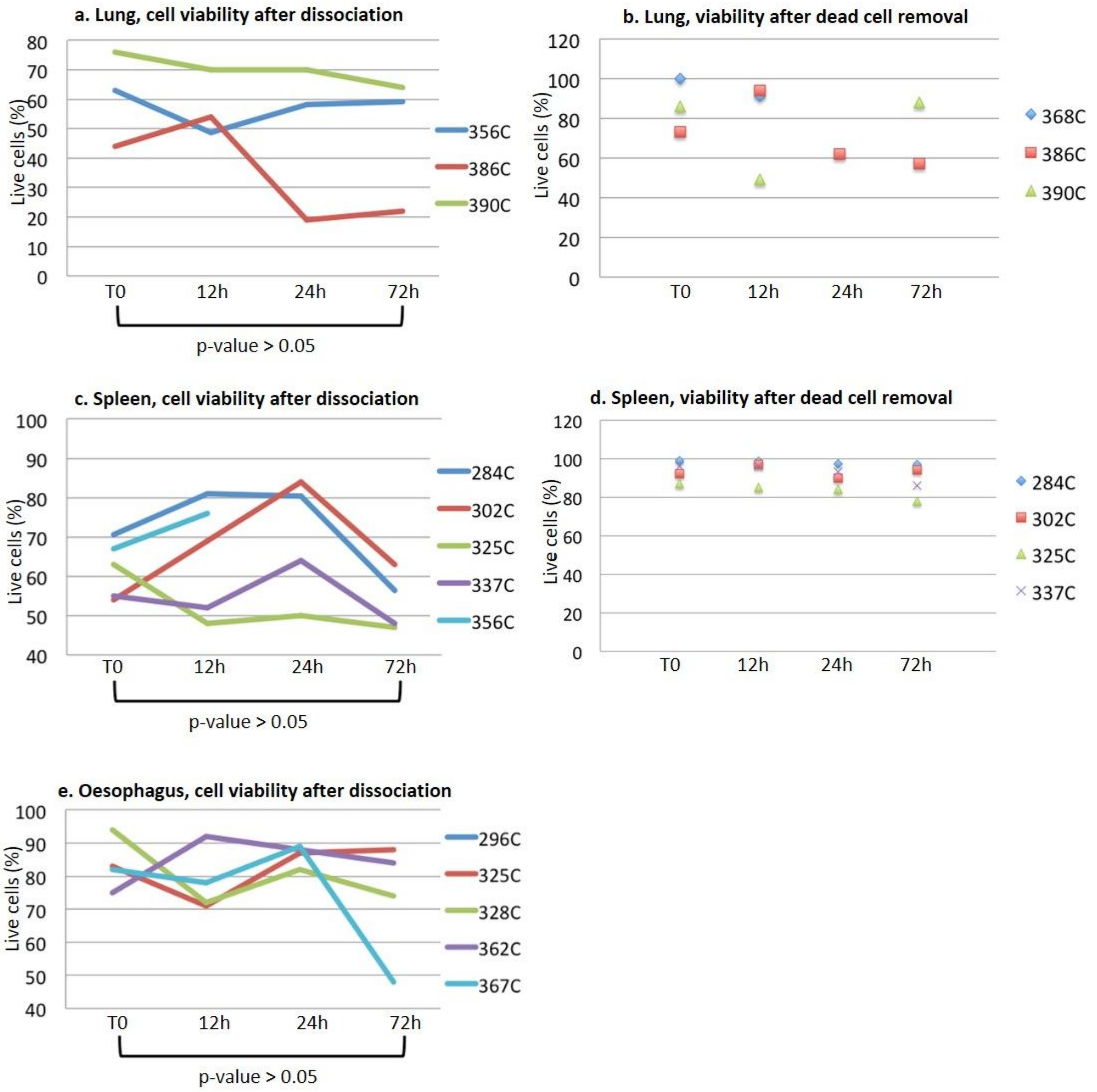


**Supplementary Figure 6. Cell viability directly after dissociation and after dead cell removal.**

Cell viabilities are shown for lung (a, b) and spleen (c, d) both directly after dissociation (a, c) and after dead cell removal (b, d), and for oesophagus only directly after dissociation (e). Student's t-tests were performed in each tissue for viability percentages directly after dissociation between T0 and 72h.

a

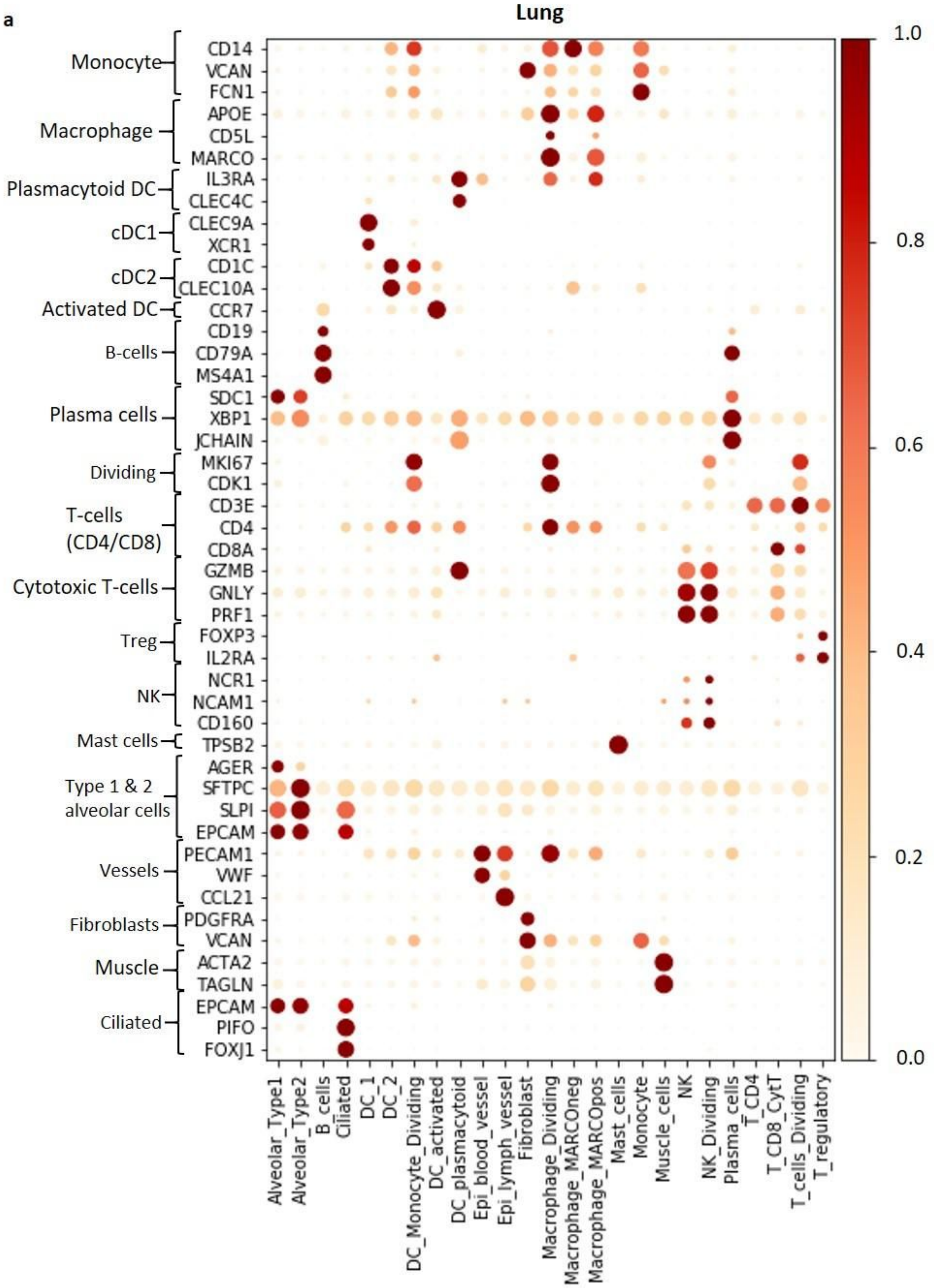
Lung
Monocyte
Macrophage
Plasmacytoid DC
cDC1
cDC2
Activated DC
B-cells
Plasma cells
Dividing
T-cells (CD4/CD8)
Cytotoxic T-cells
Treg
NK
Mast cells
Type 1 & 2 alveolar cells
Vessels
Fibroblasts
Muscle
Ciliated
CD14
VCAN
FCN1
APOE
CD5L
MARCO
IL3RA
CLEC4C
CLEC9A
XCR1
CD1C
CLEC10A
CCR7
CD19
CD79A
MS4A1
SDC1
XBP1
JCHAIN
MKI67
CDK1
CD3E
CD4
CD8A
GZMB
GNLY
PRF1
FOXP3
IL2RA
NCR1
NCAM1
CD160
TPSB2
AGER
SFTPC
SLPI
EPCAM
PECAM1
VWF
CCL21
PDGFRA
VCAN
ACTA2
TAGLN
EPCAM
PIFO
FOXJ1
Alveolar_Type1
Alveolar_Type2
B_cells
Ciliated
DC_1
DC_2
DC_Monocyte_Dividing
DC_activated
DC_plasmacytoid
Epi_blood_vessel
Epi_lymph_vessel
Fibroblast
Macrophage_Dividing
Macrophage_MARCOneg
Macrophage_MARCOpos
Mast_cells
Monocyte
Muscle_cells
NK
NK_Dividing
Plasma_cells
T_CD4
T_CD8_CytT
T_cells_Dividing
T_regulatory
1.0
0.8
0.6
0.4
0.2
0.0

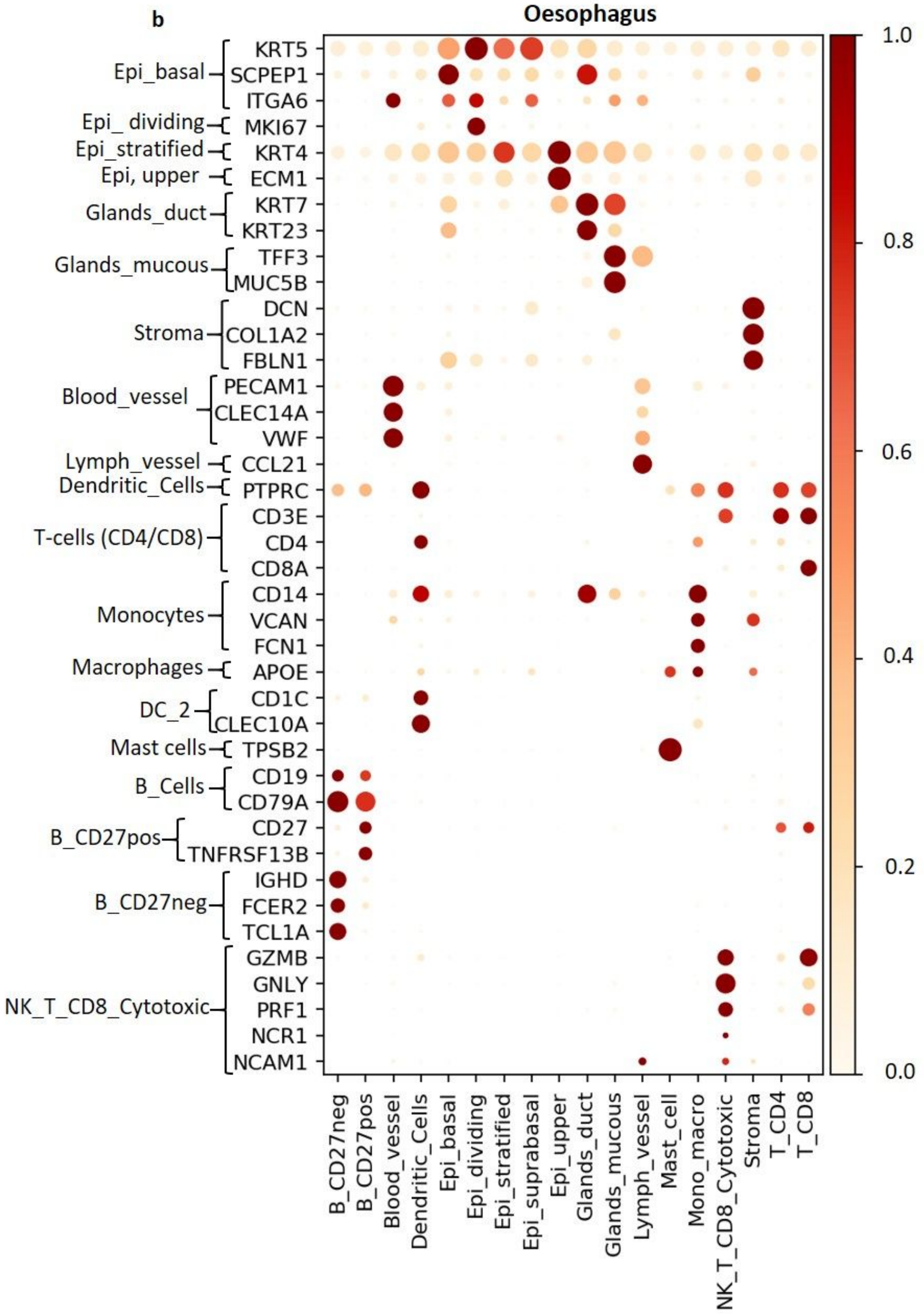
b
Oesophagus
Epi_basal
KRT5
SCPEP1
ITGA6
Epi_ dividing
MKI67
Epi_stratified
KRT4
Epi, upper
ECM1
Glands_duct
KRT7
KRT23
Glands_mucous
TFF3
MUC5B
Stroma
DCN
COL1A2
FBLN1
Blood_vessel
PECAM1
CLEC14A
VWF
Lymph_vessel
CCL21
Dendritic_Cells
PTPRC
T-cells (CD4/CD8)
CD3E
CD4
CD8A
Monocytes
CD14
VCAN
FCN1
Macrophages
APOE
DC_2
CD1C
CLEC10A
Mast cells
TPSB2
B_Cells
CD19
CD79A
B_CD27pos
CD27
TNFRSF13B
B_CD27neg
IGHD
FCER2
TCL1A
NK_T_CD8_Cytotoxic
GZMB
GNLY
PRF1
NCR1
NCAM1
B_CD27neg
B_CD27pos
Blood_vessel
Dendritic_Cells
Epi_basal
Epi_dividing
Epi_stratified
Epi_suprabasal
Epi_upper
Glands_duct
Glands_mucous
Lymph_vessel
Mast_cell
Mono_macro
NK_T_CD8_Cytotoxic
Stroma
T_CD4
T_CD8
1.0
0.8
0.6
0.4
0.2
0.0

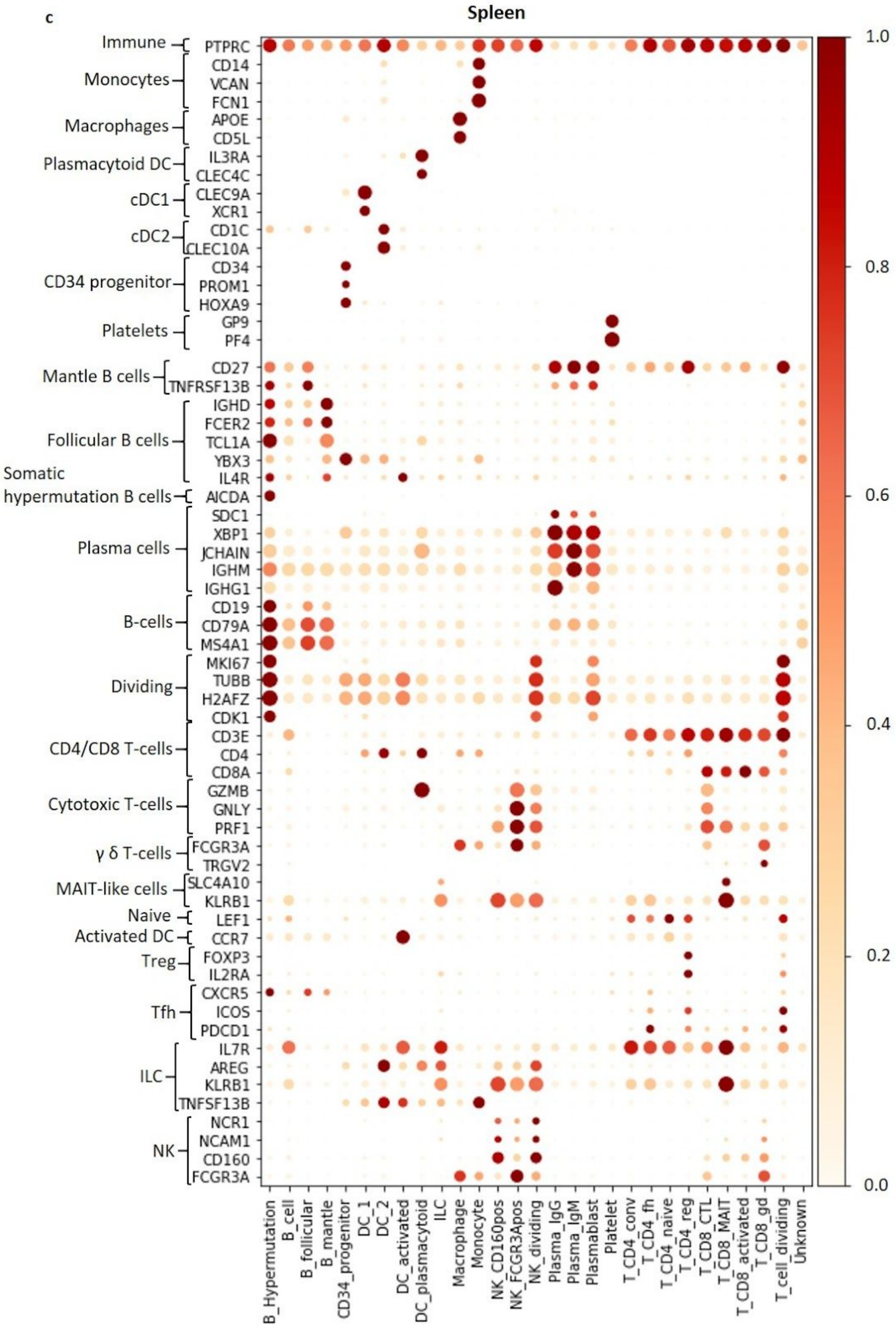


**Supplementary Figure 7. Cell type markers and their expression in the data.** Overview of Lung (a), Oesophagus (b) and Spleen (c) single-cell RNA-sequencing data cell type markers genes expression. Color represents maximum-normalised mean expression of marker genes in each cell group, and size indicates the proportion of cells expressing marker gene.

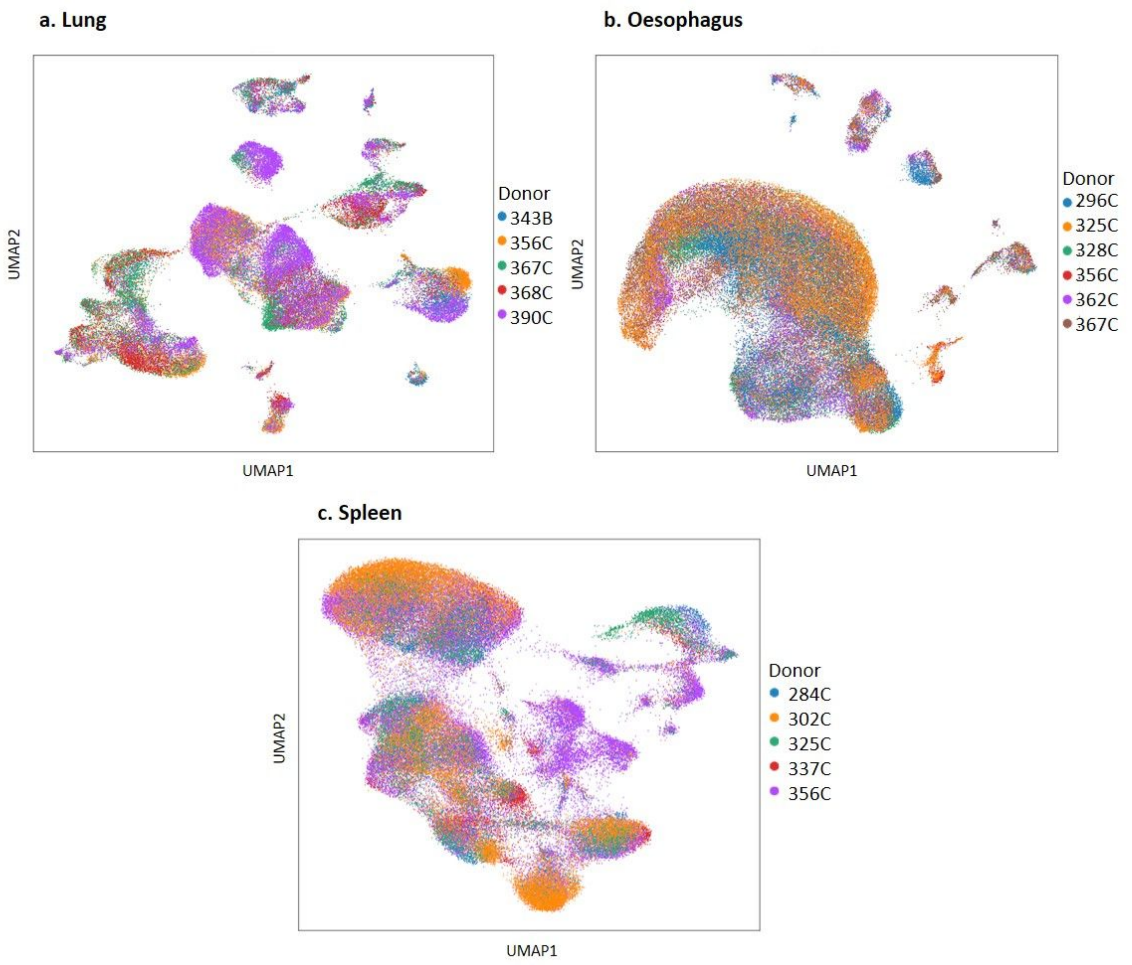


**Supplementary Figure 8: Distribution of cells from different donors.**

UMAP plots for all three organs coloured by donor, for lung (a), oesophagus (b) and spleen (c).

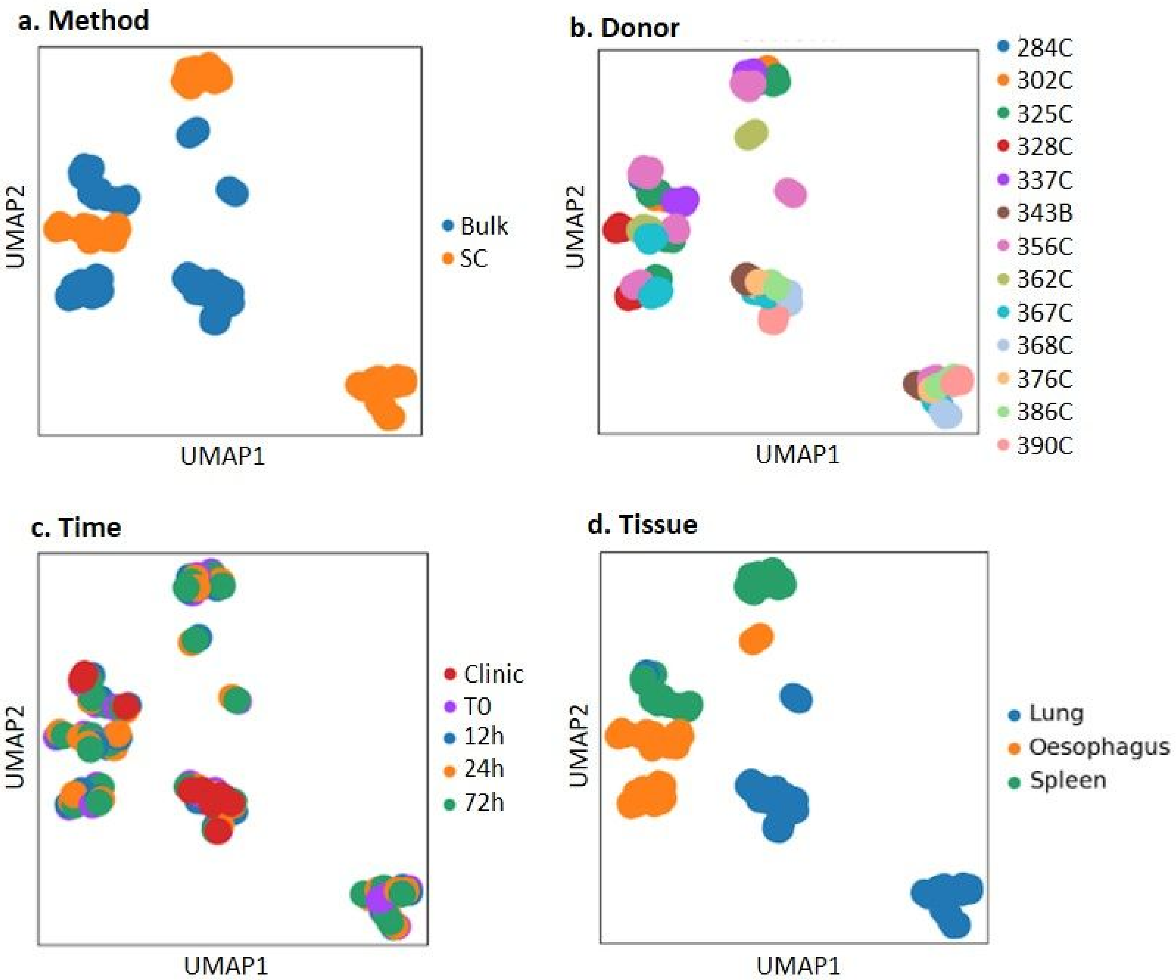


**Supplementary Figure 9. Bulk RNA-sequencing data comparison with single-cell RNA-sequencing.**

The single-cell data was combined as pseudo-bulk per sample, PCA was performed for highly variable genes across samples, UMAP coordinates were calculated based on PCA. The clusters of samples are shown on UMAP plot, colored by method (a), donor (b), time (c) and tissue (d).

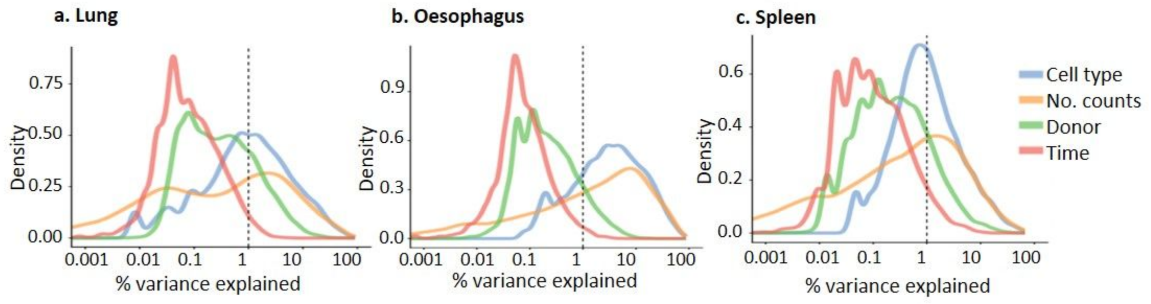


**Supplementary Figure 10. Time explains the least of variance in gene expression.**

Percentage of variance in gene expression explained by cell type, number of counts, donor and time in lung (**a**), oesophagus (**b**) and spleen (**c**). Gene-wise density plots of the gene-wise marginal $R^2$ for each variable is shown.

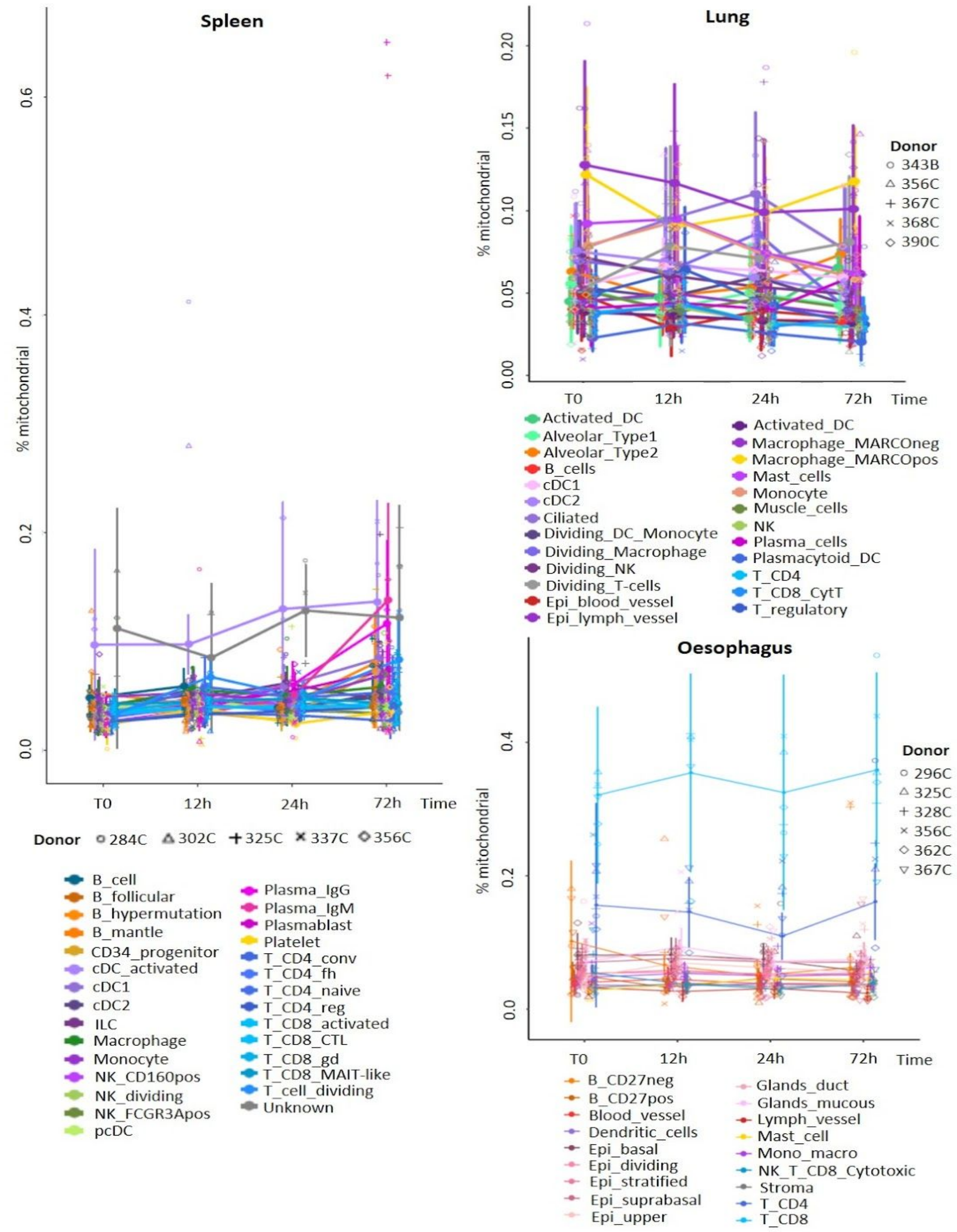


**Supplementary Figure 11. Mitochondrial percentage differs between cell types.**

Mean mitochondrial percentage and standard deviation across cells and donors was calculated per every cell type in every time point. Dots represent the mean mitochondrial percentage across donors for a cell type and time point. Whiskers show standard deviation. The same cell types are connected by line. Mean mitochondrial percentages are shown separately for every donor, cell type and time point by shapes corresponding to donors. All colors represent cell types.

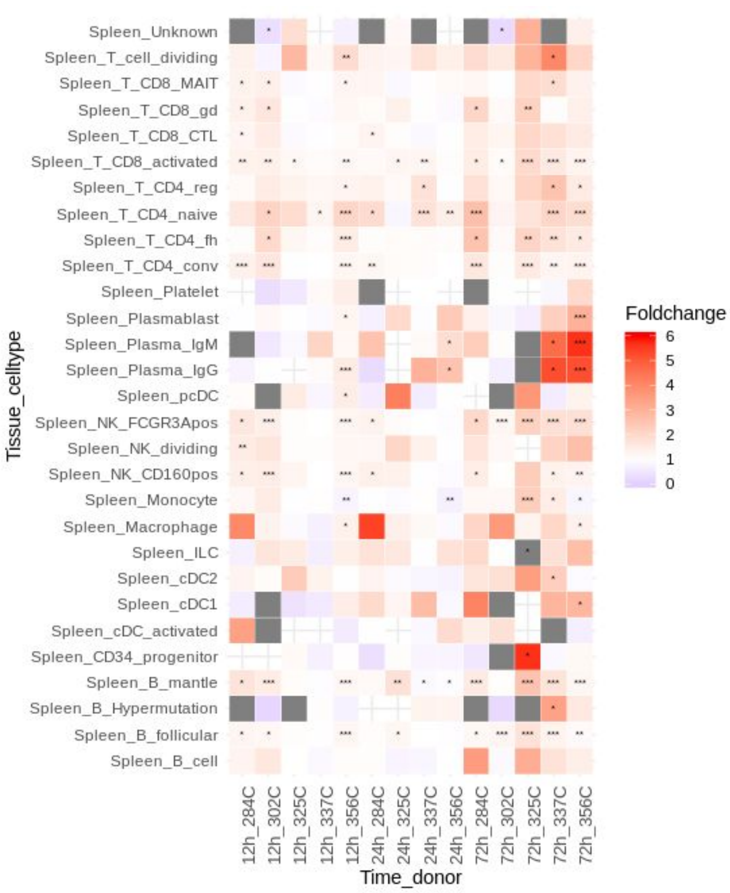


**Supplementary Figure 12. Change in the percentage of mitochondrial reads in time and by cell type and donor in spleen.**

The fold change (FC) of mitochondrial read percentage is shown for every cell type between later timepoints (12h, 24h and 72h) and time T0. FC is indicated by color with white indicating no fold change (FC=1), blue indicates a drop in mitochondrial reads compared to T0 (FC<1), red indicates increase in mitochondrial percentage compared to T0 (FC>1). Benjamin and Hochberg adjusted p-values are indicated by asterisk as follows: p-val<0.01*, p-val < 0.00001** and p-val < 0.00000001***. All cells are used, including those with high mitochondrial percentage (>10%) whose cell type annotations were derived via scmap tool based on similarity to the annotations to cells with lower mitochondrial percentage. Time points with fewer than 5 cells per time point are shown in grey. Missing values (no cells in either comparison) are shown by light grey crosses.

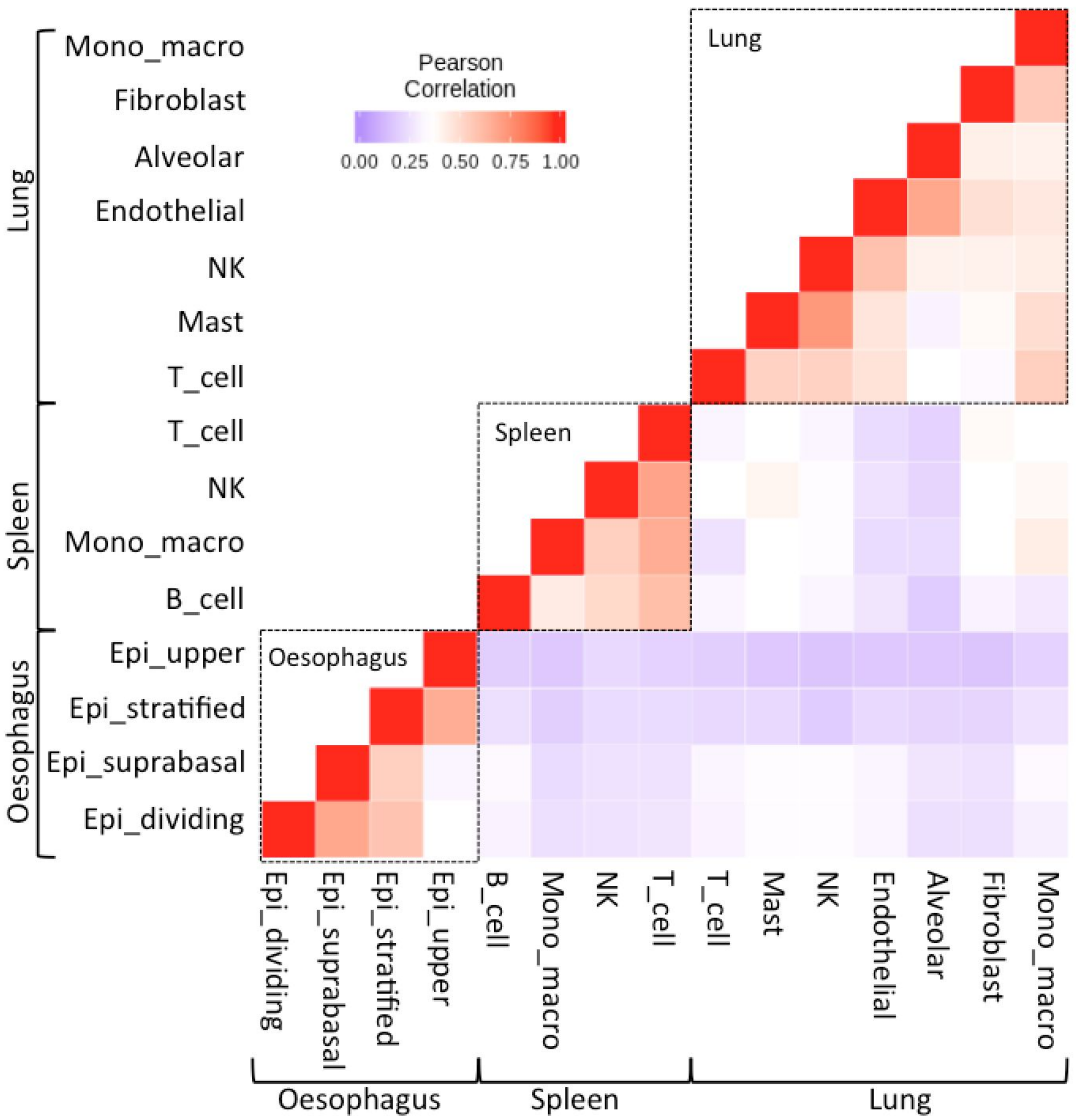


**Supplementary Figure 13. Gene signatures associated with storage time are correlated with tissue type and not cell type.**

For the major cell types for each tissue (see x axis), we calculated the explanatory variance in gene expression over time for all genes. Using this matrix we then examined the correlation between cell types and these are plotted as Pearson correlation coefficients, indicated by color.

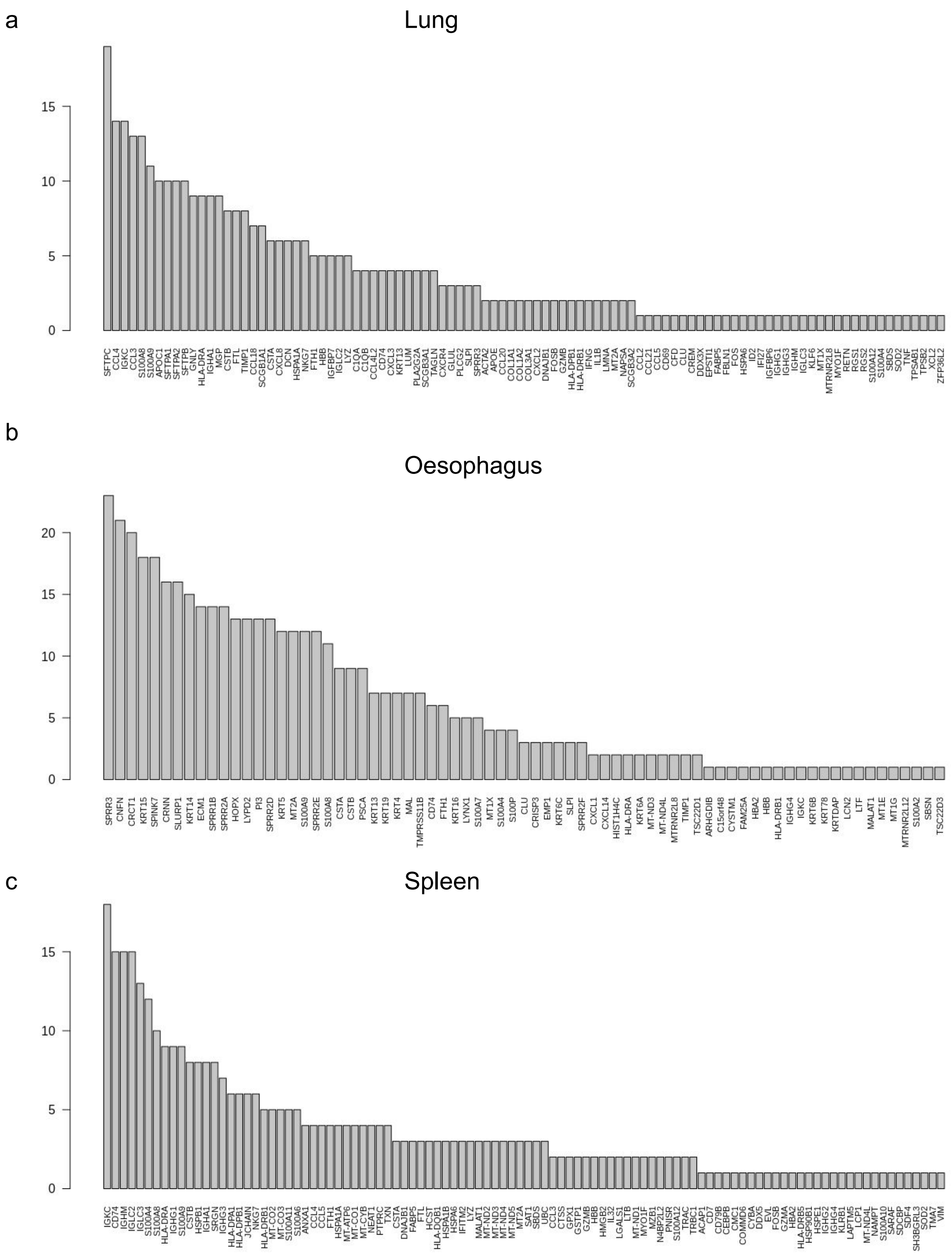


**Supplementary Figure 14**. **Frequency plots of top ambient RNA contamination genes.**

Frequency of top 20 contaminating genes from any of the samples in lung (a), oesophagus (b) and spleen (c) tissues. SoupX algorithm infers genes for a 10x run that have the likelihood for contaminating the samples. Top 20 of these genes per sample were used for calculating the frequency across samples per tissue.
